# Weak neuronal glycolysis sustains cognition and organismal fitness

**DOI:** 10.1101/2023.09.02.556016

**Authors:** Daniel Jimenez-Blasco, Jesús Agulla, Rebeca Lapresa, Marina Garcia-Macia, Veronica Bobo-Jimenez, Dario Garcia-Rodriguez, Israel Manjarrez-Raza, Emilio Fernandez, Yannick Jeanson, Spiro Khoury, Jean-Charles Portais, Daniel Padro, Pedro Ramos-Cabrer, Peter Carmeliet, Angeles Almeida, Juan P. Bolaños

## Abstract

The energy cost of neuronal activity is mainly sustained by glucose^1,2^. However, in an apparent paradox, neurons only weakly metabolize glucose through glycolysis^3,4,5,6^, a circumstance that can be accounted for by the constant degradation of 6-phosphofructo-2-kinase/fructose-2,6-bisphosphatase-3 (Pfkfb3)^3,7,8^, a key glycolysis-promoting enzyme. To evaluate the *in vivo* physiological significance of this hypo-glycolytic metabolism, here we genetically engineered mice with their neurons transformed into active glycolytic cells through Pfkfb3 expression. *In vivo* molecular, biochemical, and metabolic flux analyses of these neurons revealed an accumulation of anomalous mitochondria, complex I disassembly, bioenergetic deficiency and mitochondrial redox stress. Notably, glycolysis-mediated NAD^+^ reduction impaired sirtuin-dependent autophagy. Furthermore, these mice displayed cognitive decline and a metabolic syndrome that was mimicked by confining Pfkfb3 expression to hypothalamic neurons. Neuron-specific genetic ablation of mitochondrial redox stress corrected these alterations. Thus, the weak glycolytic nature of neurons is required to sustain higher-order organismal functions.

---

The proteolytic destabilization of Pfkfb3 in neurons takes place upon polyubiquitination by the E3 ubiquitin ligase anaphase-promoting complex/cyclosome (APC/C)^3^. APC/C can be activated by either of the cofactors Cdh1 or Cdc20^7^. However, Cdc20 is not present in differentiated neurons^8^, hence Cdh1 is the only cofactor responsible for the observed high APC/C activity that leads to Pfkfb3 protein destabilization and neuronal hypoglycolysis^3^. Stabilization of endogenous neuronal Pfkfb3 takes place in several disease conditions^9,10^ and during development^6^. In addition, aberrant hyper-glycolysis takes place in Alzheimer’s disease neurons^11^. However, the *in vivo* physiological significance of the weak adult neuronal glycolysis remains unknown, a limitation that has led to controversies^1^ thus hindering a better knowledge of brain function in health and disease. To address this issue, we aimed to generate a mouse genetic model able to boost glycolysis in neurons by means of Pfkfb3 expression during adulthood. First, conditional *Cdh1* knockout (*Cdh1^lox/lox^*) mice^12^ were mated with mice expressing Cre recombinase under the control of the neuron-specific calcium-calmodulin protein kinase-II*α* (*CamkIIα*) promoter, which is widely expressed across the brain, mainly in the hippocampus, neocortex, striatum and amygdala as from the third postnatal week^13^. As expected, the *CamkIIα-Cdh1^-/-^*progeny showed Pfkfb3 protein stabilization in neurons of the adult brain (**Fig. S1a**). However, Cyclin B1 and Rock-2, i.e. two proteins known to be APC/C-Cdh1 substrates that cause neurotoxicity^8^ and cognitive decline^14^, were also found stabilized (**Fig. S1a**). These data did not persuade us that inhibition of APC/C-Cdh1 activity would be a suitable strategy to specifically assess the *in vivo* impact of active glycolysis in neurons. To overcome this issue, we next generated, by homologous recombination in the *Rosa26* locus, a transgenic mouse harboring the full-length cDNA of *Pfkfb3* (**Fig. 1a**). A floxed (*loxP*-flanked) transcriptional *Stop* cassette was incorporated between the *Pfkfb3* cDNA and the *CAG* promoter (*Pfkfb3^lox^/+*), to eventually obtain tissue- and time-specific expression of *Pfkfb3 in vivo* (**Fig. 1a**). To ascertain the efficacy of this strategy, *Pfkfb3^lox^/+* mice were mated with *CamkIIα-Cre* mice to generate *CamkIIα-Pfkfb3* (**Fig. 1a**). *Pfkfb3^lox^/+* littermates were used as controls (WT) (**Fig. 1a**). *CamkIIα-Pfkfb3* mouse brains showed enhanced Pfkfb3 protein levels, as judged by western blotting of all areas analyzed such as cortex, hippocampus, and hypothalamus (**Fig. 1b; Fig. S1b**), as well as by hippocampal immunocytochemistry (**Fig. S1c**).

**Fig. 1.**
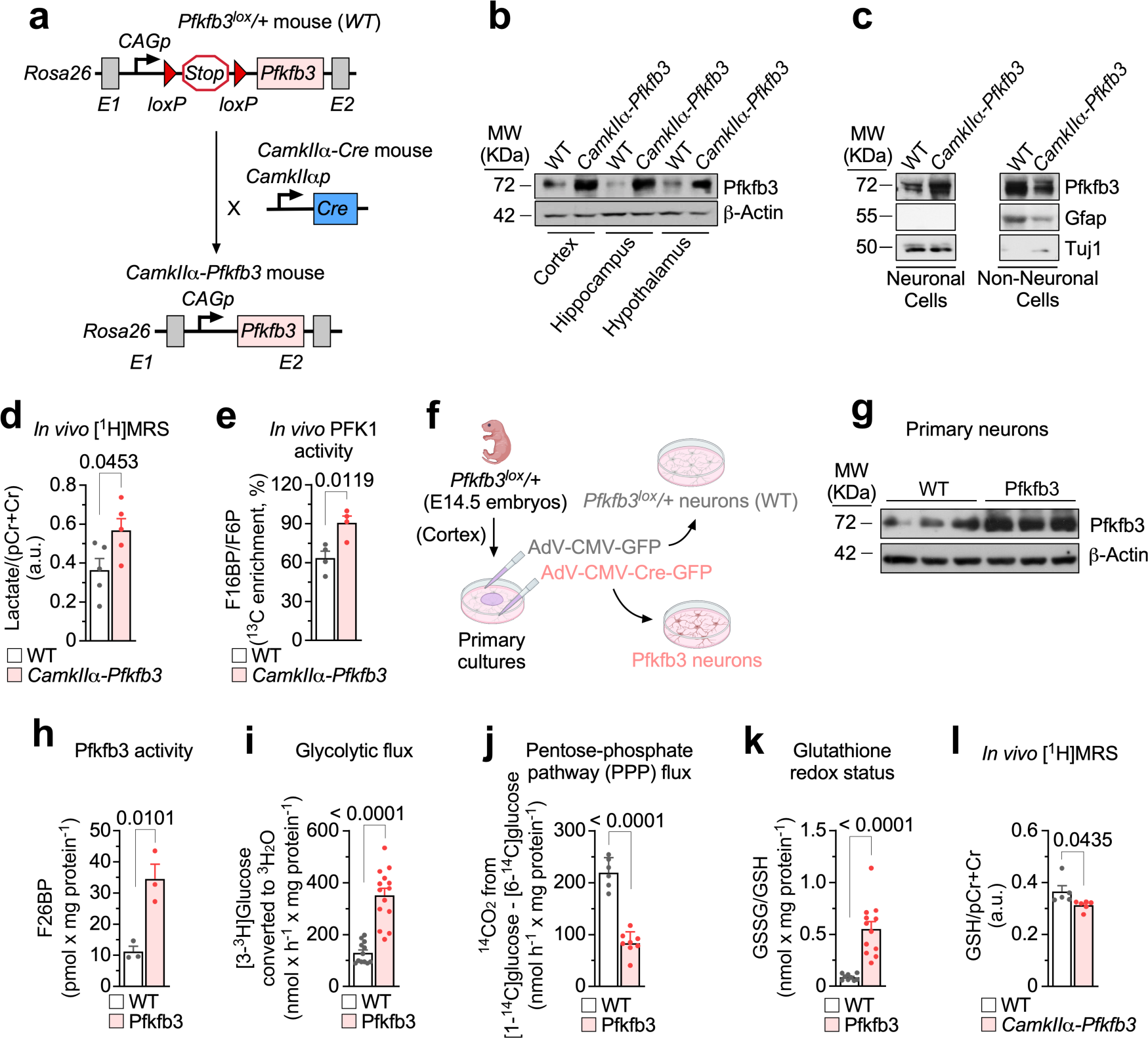
*In vivo* neuron-specific *Pfkfb3* expression activates glycolysis and inhibits PPP causing redox stress. **(a)** Strategy used to generate neuron-specific Pfkfb3-expressing (*CamkIIα-Pfkfb3*) mice. **(b)** Western blot against Pfkfb3 protein in *CamkIIα-Pfkfb3* mouse brain areas. ß-Actin was used as a loading control. (See also Figure S1b). **(c)** Western blot against Pfkfb3 protein in neurons and non-neuronal cells immunomagnetically isolated from *CamkIIα-Pfkfb3* mice. Gfap and Tuj1 were used as astrocytes or neurons enrichment and loading controls. (See also Figure S1d). **(d)** *In vivo* [^1^H]MRS analysis of lactate/(pCr+Cr) ratio in the brain of *CamkIIα-Pfkfb3* mice. Data are mean ± S.E.M. *P* value is indicated (n=5 mice per genotype; Unpaired Student’s *t*-test, two-sided). (See also Figure S1e). **(e)** *In vivo* MS analysis of F16BP/F6P ratio in the brain of *CamkIIα-Pfkfb3* mice. Data are mean ± S.E.M. *P* value is indicated (n=4 mice per genotype; Unpaired Student’s *t*-test, two-sided). (See also Figure S1f). **(f)** Adenoviral transduction strategy used to express Pfkfb3 in brain cortical neurons in primary culture from *Pfkfb3^lox^/+* mice. Created with BioRender.com. **(g)** Western blot against Pfkfb3 protein in WT or Pfkfb3 primary neurons. ß-Actin was used loading control. (See also Figure S1g). **(h-k)** F26BP concentration (h), glycolytic flux as measured by the rate of [3-^3^H]glucose conversion into ^3^H_2_O (i), PPP flux as measured by the difference in ^14^CO_2_ production from [1-^14^C]glucose and [6-^14^C]glucose (j) and glutathione redox status as measured by the ratio oxidized *versus* reduced glutathione (k) in primary neurons. Data are mean ± S.E.M. *P* value is indicated; n=3 (h), 12 (i, WT), 14 (i, Pfkfb3), 6 (j, WT), 8 (j, Pfkfb3), 9 (k, WT), 12 (k, Pfkfb3) biologically independent cell culture preparations; Unpaired Student’s *t*-test, two-sided. (See also Figures S1h, S1i). **(l)** *In vivo* [^1^H]MRS analysis of GSH/(pCr+Cr) ratio in the brain of *CamkIIα-Pfkfb3* mice. Data are mean ± S.E.M. *P* value is indicated; n=5 (WT), 6 (*CamkIIα-Pfkfb3*) mice; Unpaired Student’s *t*-test, two-sided.

In contrast to neurons, APC/C-Cdh1 activity is low in astrocytes and accounts for their naturally higher Pfkfb3 abundance and glycolytic phenotype^3^. To ascertain whether the increase in Pfkfb3 protein in the brain of the *CamkIIα-Pfkfb3* mice is neuron-specific, neurons and astrocytes were immunomagnetically purified, and the Pfkfb3 protein levels analyzed in both cell types by western blotting. As shown in **Fig. 1c (see also Fig. S1d**), the abundance of Pfkfb3 protein in *CamkIIα-Pfkfb3* neurons is like that found in wild-type astrocytes. This indicates that the stabilization of Pfkfb3 protein in neurons of the *CamkIIα-Pfkfb3* mouse model is moderately high enough to reach levels comparable with those physiologically found in neighbour astrocytes. To assess whether Pfkfb3 protein stabilization in *CamkIIα-Pfkfb3* neurons is functional, we used two strategies. First, *in vivo* ^1^H-magnetic resonance spectroscopy ([^1^H]MRS) analysis was performed in the *CamkIIα-*Pfkfb3 mouse brain (**Fig. S1e**), which revealed an enhancement in the concentration of lactate (**Fig. 1d**), suggesting increased glycolysis. And, secondly, *CamkIIα-Pfkfb3* mice were intraperitoneally injected with [U-^13^C]glucose, and glycolytic intermediates were assessed in brain extracts by mass spectrometry (MS). As shown in **Fig. 1e (**see also **Fig. S1f)**, the ratio fructose-1,6 bisphosphate (F16BP) *versus* fructose-6-phosphate (F6P) (F16BP/F6P) -i.e., the product and substrate of 6-phosphofructo-1-kinase (Pfk1), respectively-was increased, indicating Pfk1 activation in the brain of the *CamkIIα-Pfkfb3* mice. Altogether, these data indicate that the *CamkIIα-Pfkfb3* mice show efficient, otherwise moderate increase in neuronal Pfkfb3 protein that leads to Pfk1 activation and enhanced glycolytic flux *in vivo*.

To validate the glycolytic flux activation in neurons, these cells were cultured from *Pfkfb3^lox^/+* mouse embryos and, once differentiated, transduced with adenoviruses expressing Cre recombinase under the cytomegalovirus (CMV) promoter (AAV-CMV-Cre-GFP) to generate Pfkfb3 neurons. *Pfkfb3^lox^/+* neurons transduced with the AAV lacking Cre recombinase (AAV-CMV-GFP) were used as controls (WT) (**Fig. 1f**). As shown in **Fig. 1g** (see also **Fig. S1g**), Pfkfb3 protein increased, and its functional activity was confirmed as judged by the enhancement in the Pfkfb3 product, fructose-2,6-bisphosphate (F26BP) in Pfkfb3 neurons (**Fig. 1h**). These cells had higher glycolytic flux, as analyzed by the rate of [3-^3^H]glucose incorporation into ^3^H_2_O-a *bona-fide* index of glycolysis^3^ (**Fig. 1i**). Since glycolysis and pentose-phosphate pathway (PPP) are inversely correlated in neurons ^15^, we determined the PPP flux in Pfkfb3 neurons by estimating the difference in ^14^CO_2_ production from [1-^14^C]glucose-decarboxylated at the PPP and tricarboxylic acid (TCA) cycle- and [6-^14^C]glucose-decarboxylated at the tricarboxylic acid (TCA) cycle^16,17^. As shown in **Fig. 1j (**see also **Fig. S1h)**, PPP flux decreased in Pfkfb3 neurons. By producing NADPH(H^+^), PPP sustains the regeneration of reduced glutathione (GSH) from its oxidized form (GSSG)^17^. Accordingly, GSH (reduced and total forms) decreased and GSSG increased in Pfkfb3 neurons, resulting in enhanced glutathione oxidation (increased GSSG/GSH ratio) (**Fig. 1k; Fig. S1i**). These results were confirmed *in vivo* by brain [^1^H]MRS analysis in *CamkIIα-Pfkfb3* mice, as judged by the decreased GSH signal (**Fig. 1l**). These data agree with previous observations indicating that the PPP is an advantageous metabolic pathway for neurons^17–19^.

Given that GSH is an antioxidant, we next searched for possible redox stress in Pfkfb3 neurons. Reactive oxygen species (ROS) were thus assessed using two approaches. Using AmplexRed^®^ we observed an increase in fluorescence that was potentiated by rotenone (**Fig. 2a**), suggesting enhanced ROS by forward electron transfer at mitochondrial complex I. We next used the mitochondrial-specific probe MitoSox^®^, which revealed increased mitochondrial ROS (**Fig. 2b**). To further confirm the mitochondrial origin of these ROS, we used a genetic approach to express mitochondrial matrix-tagged catalase (*mCat^lox^/+*)^20^. Mating *Pfkfb3^lox^/+* with *mCat^lox^/+* generated double transgenic *Pfkfb3^lox^/+;mCat^lox^/+* mice and siblings harboring either genotype alone, namely *Pfkfb3^lox^/+* or *mCat^lox^/+* (**Fig. S2a**). Littermate embryos were individually genotyped and used to obtain neurons in primary culture (**Fig. S2b**). Transduction of *Pfkfb3^lox^/+*, *mCat^lox^/+* and *Pfkfb3^lox^/+;mCat^lox^/+* with AAV-CMV-Cre-GFP yielded Pfkfb3, mCat or Pfkfb3-mCat neurons, respectively (**Fig. S2b**); controls were *Pfkfb3^lox^/+* cells transduced with AAV-CMV-GFP (**Fig. S2b**). The increases in rotenone-induced AmplexRed® and MitoSox® fluorescence signal enhancements observed in Pfkfb3 neurons were abolished in Pfkfb3-mCat neurons (**Fig. 2a,b**), confirming the mitochondrial matrix origin of the ROS signal (mROS). To corroborate these findings *in vivo*, *CamkIIα-Pfkfb3* mice were intravenously transduced with adenoviruses expressing GFP under the control of the neuron-specific promoter hSyn (AAV-PHP.eb-hSyn-GFP) (**Fig. 2c**). Hippocampus and hypothalamus were then dissociated, and GFP^+^ cells analyzed by flow cytometry for MitoSox® fluorescence, the intensity of which was enhanced in both brain areas (**Fig. 2d**). To abolish mROS in neurons *in vivo*, the double transgenic *Pfkfb3^lox^/+;mCat^lox^/+* mice were mated with *CamkIIα-Cre* to generate *CamkIIα-Pfkfb3-mCat* mice (**Fig. S2a**). *Pfkfb3^lox^/+* (WT) and *CamkIIα-mCat* were used as controls for *CamkIIα-Pfkfb3* and *CamkIIα-Pfkfb3-mCat* mice, respectively (**Fig. S2a**). As depicted in **Fig. 2d**, the increased mROS of neurons from *CamkIIα-Pfkfb3* mice was not observed in the *CamkIIα-Pfkfb3-mCat* mice. These data indicate that glycolytically active neurons undergo mitochondrial redox stress *in vivo*.

**Fig. 2.**
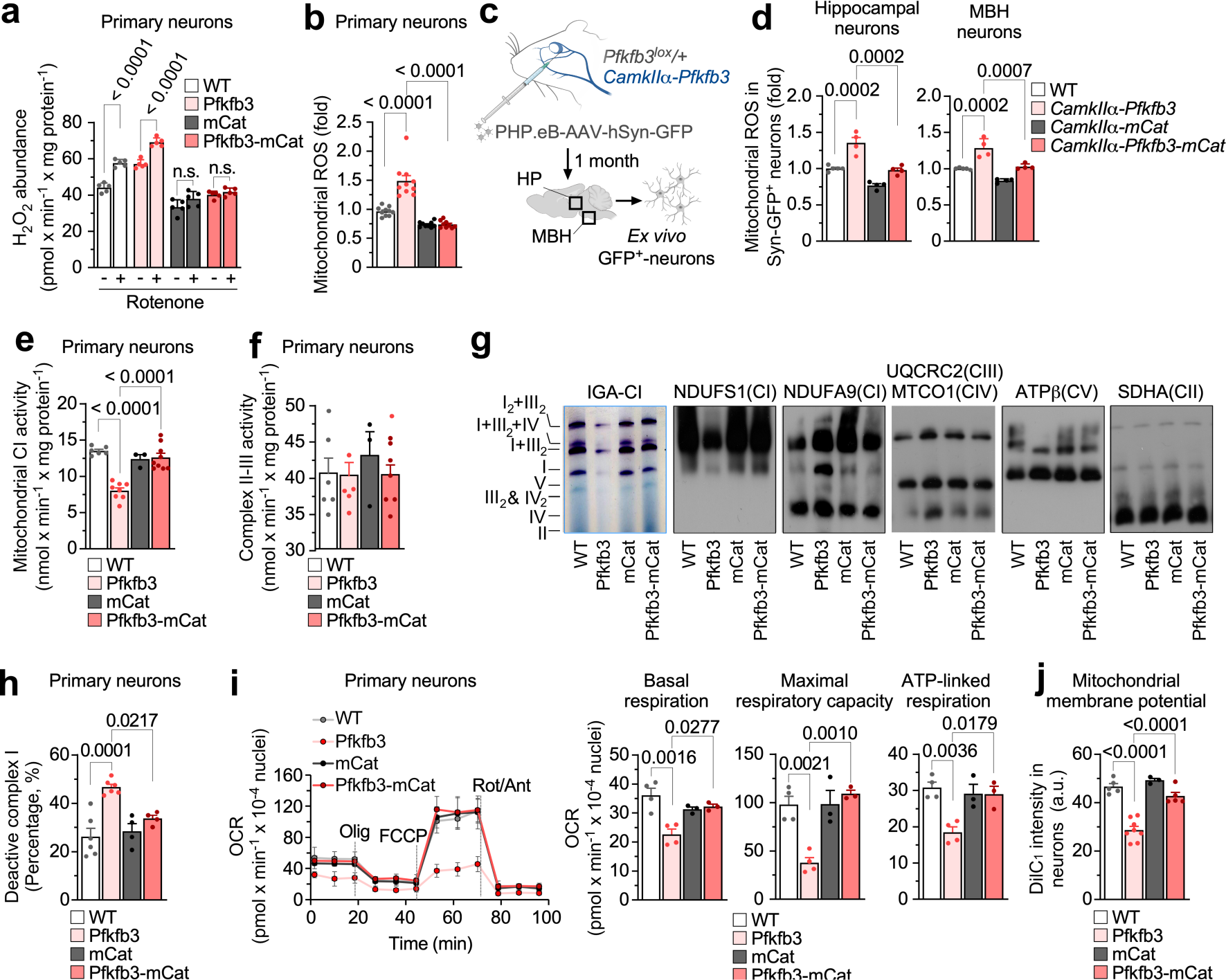
Neuron-specific *Pfkfb3* expression impairs mitochondrial bioenergetics *via* enhanced mitochondrial ROS. (a) H_2_O_2_ production in primary neurons treated or not with rotenone. Data are mean ± S.E.M. *P* value is indicated; n=5 biologically independent cell culture preparations; two-way ANOVA followed by Tukey. (b) Mitochondrial ROS production in primary neurons. Data are mean ± S.E.M. *P* value is indicated; n=10 (WT, Pfkfb3) or 9 (mCat, Pfkfb3-mCat) biologically independent cell culture preparations; one-way ANOVA followed by Tukey. (c) *In vivo* adeno-associated viral intravenous transduction strategy to express GFP in the neurons of *Pfkfb3^lox^/+* or *CamkIIα-Pfkfb3* mice. Created with BioRender.com. (See also Figure S2f). (d) Mitochondrial ROS production in hippocampal or MBH neurons freshly isolated from mice of the different genotypes previously transduced with adeno-associated viral particles expressing GFP under the neuron-specific *hSyn* promoter. Data are mean ± S.E.M. *P* value is indicated; n=5 (WT) or 4 (*CamkIIα-Pfkfb3*, *CamkIIα-mCat, CamkIIα-Pfkfb3-mCat*) mice; one-way ANOVA followed by Bonferroni. (e) Mitochondrial complex I activity in primary neurons. Data are mean ± S.E.M. *P* value is indicated; n=6 (WT), 8 (Pfkfb3), 3 (mCat) or 9 (Pfkfb3-mCat) biologically independent cell culture preparations; one-way ANOVA followed by Bonferroni. (f) Mitochondrial complex II-III activity in primary neurons. Data are mean ± S.E.M. *P* value is indicated; n=6 (WT, Pfkfb3), 3 (mCat) or 7 (Pfkfb3-mCat) biologically independent cell culture preparations;one-way ANOVA followed by Bonferroni. (g) In-gel activity of complex I (IGA-CI) and blue-native gel electrophoresis (BNGE) followed by immunoblotting against CI subunits NDUFS1 and NDUFA9, CIII subunit UQCRC2, CIV subunit MTCO1, CV subunit ATPß or CII subunit SDHA in primary neurons. (See also Figure S2c). (h) Deactive mitochondrial complex I activity in primary neurons. Data are mean ± S.E.M. *P* value is indicated; n=6 (WT, Pfkfb3) or 4 (mCat, Pfkfb3-mCat) biologically independent cell culture preparations; one-way ANOVA followed by Bonferroni. (i) Oxygen consumption rates (OCR) analysis and calculated parameters in primary neurons. Data are mean ± S.E.M. *P* value is indicated; n=4 (WT, Pfkfb3) or 3 (mCat, Pfkfb3-mCat) biologically independent cell culture preparations; one-way ANOVA followed by Bonferroni. (j) Mitochondrial membrane potential (ΔY_m_) in primary neurons. Data are mean ± S.E.M. *P* value is indicated; n=5 (WT), 8 (Pfkfb3), 3 (mCat) or 5 (Pfkfb3-mCat) biologically independent cell culture preparations; one-way ANOVA followed by Bonferroni. (See also Figure S2d, S2f).

Since excess mROS may cause mitochondrial damage^21^, we next assessed the mitochondrial respiratory chain (MRC) in Pfkfb3 neurons. As sown in **Fig. 2e**, mitochondrial complex I (CI) activity was impaired in Pfkfb3, but not in Pfkfb3-mCat neurons; however, complex II-III was unaffected (**Fig. 2f**). To further explore this, we determined the MCR assembly of mitochondrial proteins by blue-native gel electrophoresis (BNGE). Pfkfb3 neurons showed a selective loss of free CI, according to the in-gel activity (IGA) assay (**Fig. 2g**, **Fig. S2c**). CI specific activity analysis in Pfkfb3 neurons revealed an increase in the deactive^22^ form of the complex, which was not observed in the Pfkfb3-mCat cells (**Fig. 2h**), indicating that mROS mediates CI deactivation and inhibition. Notably, CI lost its N-(NADH-binding) module exclusively in its free form, not in that assembled in supercomplexes (SCs), whereas its Q-(ubiquinone-binding) module was intact (**Fig. 2g**, **Fig. S2c**). Since these data suggest MRC dysfunction, we next performed a bioenergetic profile analysis using the Seahorse® technology. This revealed an impairment in basal, maximal, and ATP-linked oxygen consumption rates (OCR) in Pfkfb3 neurons that was corrected in Pfkfb3-mCat (**Fig. 2i**). Flow cytometry (**Fig. 2i**) and confocal microscopy (**Fig. S2d**) analyses manifested loss in the inner mitochondrial membrane potential (ΔΨ_m_) in Pfkfb3 neurons, an effect that was canceled by mCat. Metabolomics analysis in the brain of the *CamkIIα-Pfkfb3* mice revealed significant alterations in metabolic pathways, including a reduced concentration of the TCA cycle intermediary citrate (**Fig. S2e**). To confirm bioenergetic failure of neurons *in vivo*, hippocampal (HP) and mediobasal hypothalamic (MBH) GFP^+^ cells (**Fig. 2c**) were analyzed by flow cytometry, which showed ΔΨ_m_ loss in the *CamkIIα-Pfkfb3* mice that was corrected in *CamkIIα-Pfkfb3-mCat* (**Fig. S2f**). These data indicate that neurons with active glycolysis undergo mROS-mediated deactivation of mitochondrial CI causing a bioenergetic failure.

Cytosolic NAD^+^ regeneration is required for glycolysis^23,24^. However, dysfunctional mitochondria, such as that taking place in Pfkfb3 neurons, have impaired their ability to regenerate NAD^+24,25^. Determination of the rates of glucose consumption and lactate production increased by ∼2.5-fold and 1.3-fold, respectively (**Fig. S3a**), showing that pyruvate to lactate conversion, whilst discretely increased by Pfkfb3 expression (see, also, **Fig. 1d** *in vivo*), does not wholly account for the high increase in the rate of glucose consumption. This is in good agreement with the occurrence in neurons of lactate dehydrogenase isoform-1 (LDH1)^4,26^, with low pyruvate-to-lactate conversion rate and, therefore, weak capacity to regenerate NAD^+^. We then sought to ascertain whether glucose would be transformed into lipids, a process that requires the conversion of pyruvate into citrate, which leaves mitochondria for lipid synthesis^27,28^. Interestingly, MS analysis of brain extracts from *CamkIIα-Pfkfb3* mice showed increased [U-^13^C]glucose conversion into [^13^C]lipids (**Fig. S3b**). In addition, the metabolomics analysis in the brain of these mice revealed data compatible with reduced lipolysis (**Fig. S2h**). Finally, [U-^14^C]glucose conversion into [^14^C]lipids in Pfkfb3 neurons increased (**Fig. 3a**) at the same rate of glycolysis (**Fig. 1h**), suggesting that the vast majority of glycolytically-consumed glucose is destined for lipids. Since lipogenesis from glucose is not an NAD^+^ regenerating process, we assessed NAD^+^ levels. As shown in **Fig. 3b**, NAD^+^ concentration was decreased in Pfkfb3 neurons, suggesting that active glycolysis causes net NAD^+^ consumption. This was confirmed by incubating Pfkfb3 neurons with the NAD^+^-precursor, nicotinamide mononucleotide (NMN), which fully restored NAD^+^ levels (**Fig. 3b**). NAD^+^ is an essential substrate for sirtuin deacetylase activity^29^, which therefore should be affected by NAD^+^ loss. Although histone deacetylase (HDAC) activity - reflecting total sirtuin activity-was unaltered (**Fig. 3c**), sirtuin-specific activity was impaired in Pfkfb3 neurons, an effect that was rescued by NMN (**Fig. 3d**). Interestingly, Atg7, a protein that requires sirtuin-mediated deacetylation to promote autophagy^29^, was hyperacetylated (**Fig. 3e**, **Fig. S3c**) suggesting impaired autophagy in glycolytically active neurons. To test this end, we determined the autophagic flux, which showed to be decreased in Pfkfb3 neurons, an effect that was rescued by NMN (**Fig. 3f**, **Fig. S3d**). This end was confirmed *ex vivo* in the cortex and hippocampus freshly dissected from the *CamkIIα-Pfkfb3* brain, where both the rates of autophagy and mitophagy were reduced (**Fig. 3g**, **Fig. S3e**). These data indicate that glycolytically active neurons show an impaired autophagic flux and that, by modulating the NAD^+^ pool, glycolysis regulates autophagy.

**Fig. 3.**
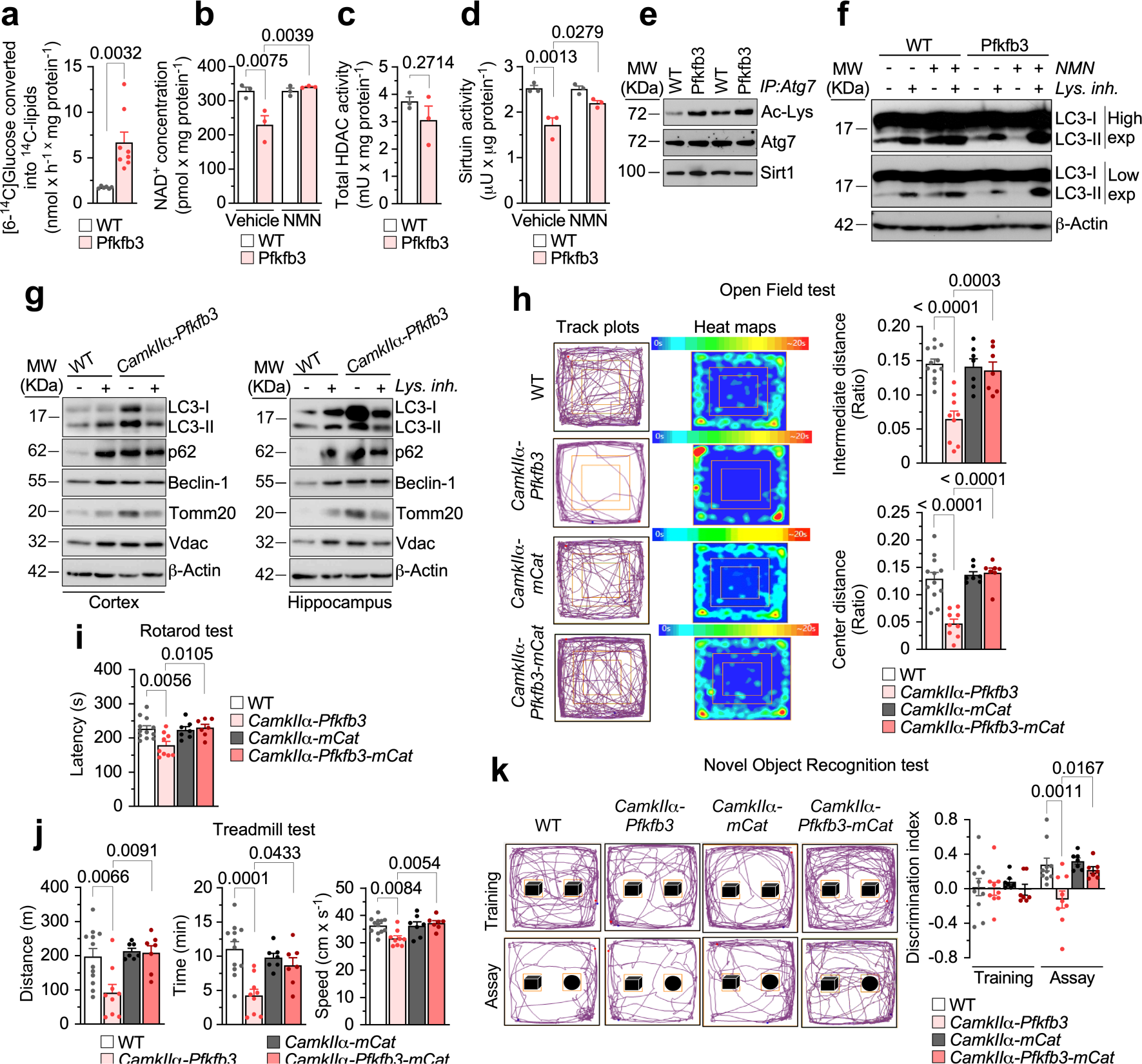
Neuron-specific *Pfkfb3* expression inhibits sirtuin-mediated autophagy causing motor and cognitive impairments *via* mitochondrial ROS. **(a-d)** Lipogenesis rate as measured by the rate of [6-^14^C]glucose conversion into ^14^C-lipids (a), NAD^+^ concentration (b), total HDAC activity (c) and sirtuin activity (d) in primary neurons. Data are mean ± S.E.M. *P* value is indicated; n=6 (a, WT), 8 (a, Pfkfb3) or 3 (b, c, d) biologically independent cell culture preparations; Unpaired Student’s *t*-test, two-sided (a, c);two-way ANOVA followed by Tukey (b, d). **(e)** Immunoprecipitation of Atg7 followed by western blot against Ac-Lys, Atg7 and Sirt1 in primary neurons; n=2 biologically independent cell culture preparations are shown. (See also Figure S3c). **(f)** Autophagic flux analysis by western blot against LC3-I and LC3-II in primary neurons incubated, or not, with NMN or lysosomal inhibitors. ß-Actin was used as loading control. (See also Figure S3d). **(g)** Autophagic flux analysis by western blot against LC3-I, LC3-II, p62, Beclin-1, Tomm20 and Vdac in freshly obtained cortical and hippocampal slices from WT or *CamkIIα-Pfkfb3* mice, incubated, or not, with lysosomal inhibitors. ß-Actin was used as loading control. (See also Figure S3e). **(h-k)** Open field (h), rotarod (i), treadmill (j) and novel object recognition (k) tests in mice of the indicated genotypes. Data are mean ± S.E.M. *P* value is indicated; n=12 (WT), 9 (*CamkIIα-Pfkfb3*) or 7 (*CamkIIα-mCat, CamkIIα-Pfkfb3-mCat*) mice; one-way ANOVA followed by Bonferroni (h-j) or two-way ANOVA followed by Tukey (k). (See also Figures S3j-n).

Next, we aimed to investigate whether the metabolic alterations occurring in glycolytically active neurons impacted on their function and viability. According to the brain [^1^H]MRS analysis (**Fig. S3f**), the neuronal functional markers N-acetyl-aspartate plus its glutamyl-derivative N-acetylaspartylglutamate (NAA + NAAG) were decreased in *CamkIIα-Pfkfb3* mice, which also showed astrogliosis according to GFAP staining (**Fig. S3g**). Moreover, flow cytometry analysis in freshly isolated neurons from hippocampus and hypothalamus from *CamkIIα-Pfkfb3* mice previously transduced with AAV-PHP.eb-hSyn-GFP, revealed increased apoptotic neurons (**Fig. S3h**). Moreover, immunocytochemical analysis of the *CamkIIα-Pfkfb3* hippocampus showed disruption of neuronal integrity, according to Tuj-1 staining (**Fig. S3i1**). Notably, these alterations were corrected in the *CamkIIα-Pfkfb3-mCat* mice (**Fig. S3h,i**). These set of data *in vivo* confirm that glycolytic neurons undergo mitochondrial redox stress that contributes to neuronal dysfunction and damage. To address whether the metabolic and functional alterations that take place in glycolytically active neurons impact the organismal level, we performed a battery of behavioral tests. The open field test (**Fig. 3h; Fig. S3j**) revealed fear/anxiety in the *CamkIIα-Pfkfb3* mice. These mice performed worse at the rotarod (**Fig. 3i**) and treadmill (**Fig. 3j**), indicating loss of motor activity, poor endurance and slowness. Furthermore, the novel object recognition test showed short-term cognitive impairment (**Fig. 3k**). Female mice showed identical phenotype (**Fig. S3k-n**). Notably, these alterations were abolished in the *CamkIIα-Pfkfb3-mCat* male mice (**Fig. 3h-k; Fig. S3j**). Thus, the transformation of neurons into glycolytically active cells develops functional alterations in neurons and loss of cognitive performance.

To further characterize glycolytically active-neuron mice, we monitored weight. *CamkIIα-Pfkfb3* mice showed no weight change at 2.5 months, but it increased at the age of 8 months (**Fig. S4a**). To boost this phenotype, mice were fed with a high-fat diet (HFD) from weanling for 21 weeks. As shown in **Fig. 4a** and **Fig. S4b**, the weight of *CamkIIa-Pfkfb3* mice progressively increased, along with food intake (**Fig. 4b**), effects that were abolished in *CamkIIα-Pfkfb3-mCat* mice. In the *Cln7^Δex^*^2^ mouse model of Neuronal Ceroid Lipofuscinosis, we previously observed neuronal Pfkfb3 protein stabilization ^10^. Here, we show that chronic (2.5 months, daily) intracerebroventricular administration of the Pfkfb3-specific inhibitor AZ67^30^ in *Cln7^Δex^*^2^ mice prevented the increase in weight gain observed in vehicle-treated *Cln7^Δex^*^2^ mice (**Fig. S4c**). Subcutaneous and abdominal white fat, and brown fat (**Fig. S4d**) were increased in *CamkIIα-Pfkfb3* mice, but not in the *CamkIIα-Pfkfb3-mCat* animals. Likewise, these mice revealed hepatomegaly and sarcopenia (**Fig. S4e**). Plasma triglycerides (**Fig. 4c**), free fatty acids (**Fig. 4d**), leptin (**Fig. 4e**), and insulin (**Fig. 4f**) were enhanced in *CamkIIα-Pfkfb3* and abolished in *CamkIIα-Pfkfb3-mCat* mice. Finally, *CamkIIα-Pfkfb3* mice were unable to reduce plasma glucose levels in the glucose overload tolerance test (**Fig. 4g; Fig. S4f**), whereas *CamkIIα-Pfkfb3-mCat* mice correctly performed this test. Given that these observations suggest metabolic-like syndrome, we assessed the autophagy capacity of the mediobasal hypothalamus (MBH), as this process has been previously reported to be specifically impaired in the MBH^31^. Analysis of the MBH freshly dissected from the *CamkIIα-Pfkfb3* mice revealed impaired autophagic flux (**Fig. 4h**, **Fig. S4g**), an effect that was ablated in *CamkIIα-Pfkfb3-mCat* mice. To assess whether the anatomical origin of this phenotype is the MBH, or it is the indirect consequence of an impaired neuronal circuitry from other brain areas, we confined Pfkfb3 expression in the MBH neurons. Thus, *Pfkfb3^lox/+^*, mice were stereotaxically injected with adenovirus harboring Cre recombinase governed by the *CamkIIα* promoter (AAV-CamkIIα-Cre-GFP) in the arcuate nucleus of the MBH in order to induce Pfkfb3 expression in the neurons of this brain area (MBH-nPfkfb3) (**Fig. 4i**). At the age of 21 weeks, no alterations in behavioral tests such as the open field (**Fig. S4h**) and the novel object recognition (**Fig. S4i**) were observed, indicating lack of fear/anxiety, motor discoordination or cognitive disturbances. However, weight (**Fig. 4j, Fig. S4j**) and food intake (**Fig. 4k**) increased in MBH-nPfkfb3 mice when compared with mice injected with AAV lacking Cre recombinase (AAV-CamkIIα-GFP). Moreover, these effects were accompanied by increased weight in the white and brown (**Fig. S4k**) adipose tissue, hepatomegaly (**Fig. S4l**), and glucose intolerance (**Fig. 4l; Fig. S4m**). Altogether, these results suggest that glycolytically active neurons of the arcuate nucleus impair hypothalamic control of food intake and organismal metabolism.

**Fig. 4.**
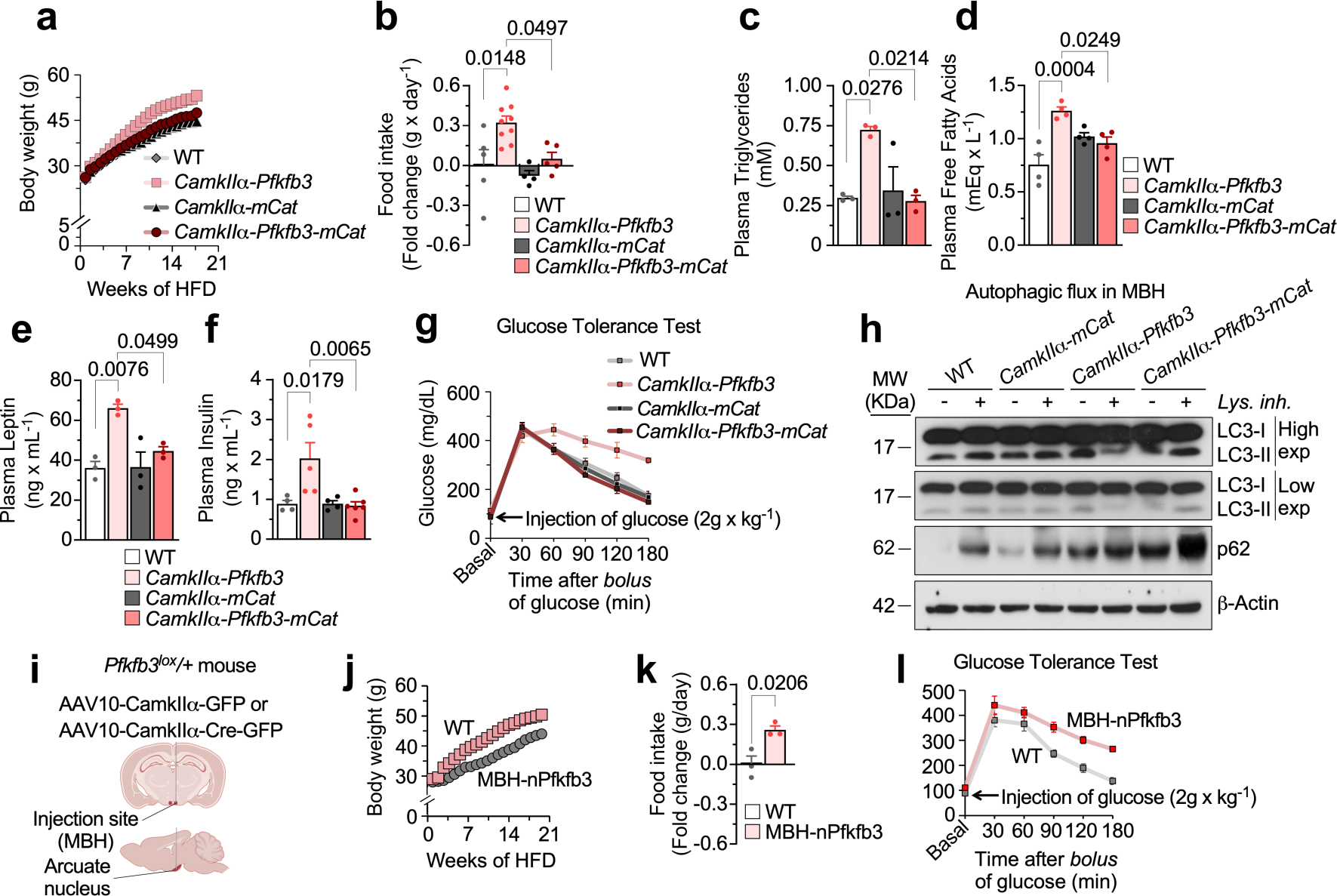
Neuronal *Pfkfb3* induces a mROS-mediated metabolic-like syndrome that is mimicked by confining Pfkfb3 expression in mediobasal hypothalamic neurons. **(a)** Body weight progressions in mice of the indicated genotypes fed with HFD. (See also Figure S4b). **(b-f)** Food intake (b), plasma triglycerides (c), free fatty acids (d), leptin (e) and insulin (f) in HFD-fed mice of the indicated genotypes. Data are mean ± S.E.M. *P* value is indicated; n=5 (b), 3 (c, e) or 4 (d, f) for WT; n= 9 (b), 3 (c, e) or 4 (d, f) for *CamkIIα-Pfkfb3*); n= 4 (b), 3 (c, e) or 4 (d, f) for *CamkIIα-mCat*); n= 5 (b), 3 (c, e), 4 (d) or 6 (f) for *CamkIIα-Pfkfb3-mCat* mice; one-way ANOVA followed by Bonferroni. **(g)** Glucose tolerance test in HFD-fed mice of the indicated genotypes. (See also Figure S4f). **(h)** Autophagic flux analysis by western blot against LC3-I, LC3-II and p62 in freshly obtained mediobasal hypothalamic (MBH) slices from mice of the indicated genotypes, incubated, or not, with lysosomal inhibitors. ß-Actin was used as loading control. (See also Figure S4g) **(i)** Strategy used to express Pfkfb3 in MBH neurons by stereotaxical injections of adeno-associated virus expressing, or not, Cre recombinase. Created with BioRender.com. **(j)** Body weight progressions in WT or MBH neurons Pfkfb3-expressing (MBH-nPfkfb3) mice fed with HFD. (See also Figure S4j) **(k)** Food intake in WT or MBH-nPfkfb3 mice fed with HFD. Data are mean ± S.E.M. *P* value is indicated (n=3 mice per condition; Unpaired Student’s *t*-test, two-sided). **(l)** Glucose tolerance test in WT or MBH-nPfkfb3 mice fed with HFD. (See also Figure S4m)

In conclusion, here we show that neurons actively preserve a hypo-glycolytic metabolism to boost pentose-phosphate pathway, therefore avoiding redox stress-mediated mitochondrial impairment. Given the inefficiency of LDH1^4,26^, functional mitochondria turn essential for neurons to regenerate cytosolic NAD^+32^ *via* the malate and glycerol phosphate shuttles^33^. Thus, the impaired mitochondria of glycolytically active neurons hinder their ability to avoid NAD^+^ depletion, causing autophagy impairment. Neurons are thus metabolically inflexible cells unable to adapt to sustained active glycolysis, unlike their neighbor astrocytes ^34^, which explains the aberrant neuronal glycolysis found in neurodegeneration^9,10,11^ and metabolic-like syndrome where neuron-specific inhibition of Pfkfb3 may be considered as a therapeutic opportunity.

## Data Availability

Source data are provided with this paper. The datasets generated and analyzed during the current study are available from the corresponding author on reasonable request.

## Acknowledgements

We acknowledge the technical assistance of Monica Resch, Monica Carabias-Carrasco, Lucia Martin and Estefania Prieto-Garcia, from the University of Salamanca. Staff members (Amandine Rocher, Hanna Kulyk, Nina Lager-Lachaud, Lara Gales, Floriant Bellvert) of MetaToul (Toulouse, France, www.metatoul.fr) are gratefully acknowledged for technical support and access to mass spectrometry facilities. This work was funded by the European Regional Development Fund, Agencia Estatal de Investigación (PID2019-105699RB-I00 / AEI / 10.13039/501100011033 and RED2018-102576-T to JPB; SAF2017-90794-REDT to AA), Instituto de Salud Carlos III (CB16/10/00282 to JPB; PI18/00285; RD16/0019/0018 to AA), Junta de Castilla y León (CS/151P20 and Escalera de Excelencia CLU-2017-03 to JPB and AA). MetaToul is part of the national infrastructure MetaboHUB, and is funded by the French National Research Agency (ANR) with grant number MetaboHUB-ANR-11-INBS-0010.

## Authors contributions

Conceived the idea and designed research: JPB Performed research: DJB, JA, RL, MGM, VBJ, DRG, DIMR, EF, YJ, SK, JCP, DP, PRC, AA Analyzed data: JPB, AA, DJB, JA, RL, MGM, VBJ, DRG, DIMR, EF, YJ, SK, JCP, DP, PRC, AA Contributed materials: PC Wrote the manuscript: JPB Edited and approved the manuscript: All co-authors

## Methods

### Mice

All protocols were performed according to the European Union Directive 86/609/EEC and Recommendation 2007/526/EC, regarding the protection of animals used for experimental and other scientific purposes, enforced in Spanish legislation under the law 6/2013. Protocols were approved by the Bioethics Committee of the University of Salamanca. All mice used in this study were of the C57Bl/6J background. Both male and female mice were used, although most data shown correspond to male mice unless otherwise stated. Mice were bred at the Animal Experimentation Facility of the University of Salamanca in cages (maximum 5 animals/cage), and a light-dark cycle was maintained for 12 hours. Humidity was 45-65% and temperature 20-25°C. Animals were fed *ad libitum* with a standard solid diet (Envigo-Harlan Tekland Global 18% Protein Rodent Diet, USA; 18% proteins, 3% lipids, 58.7% carbohydrates component, 4.3% cellulose, 4% minerals, and 12% humidity) and free access to water. When indicated, animals were switched to a high-fat diet (HFD, D12451, Research Diets, NJ, USA; 20% proteins, 45% lipids, and 35% carbohydrates, plus minerals and vitamins).

### *In vivo* generation of neuron-specific *Cdh1^-/-^* mice

To inactivate *Cdh1* gene in neurons of the adult brain, *Cdh1^lox/lox^* mice^12^ were mated with mice carrying the gene encoding *Cre* recombinase under the control of the *CamkIIα* promoter (The Jackson Laboratory)^13^, generating *CamkIIα-Cdh1^-/-^* mice.

### *In vivo* generation of neuron-specific *Pfkfb3*-expressing mice

To do this, *Pfkfb3^lox^/+* mice were generated by homologous recombination in the *Rosa26* locus of embryonic stem cells under a C57BL/6J background, where we introduced the full-length cDNA of mouse *Pfkfb3* preceded by a transcriptional STOP cassette flanked by two *loxP* sites. This loxP-flanked STOP signal was incorporated between the *CAG* promoter and the mouse *Pfkfb3* cDNA (**Fig. 1a**). To express Pfkfb3 in neurons *in vivo*, *Pfkfb3^lox^/+* mice were mated with *CamkIIα-Cre* mice to generate *CamkIIα-Pfkfb3*.

### *In vivo* generation of neuron-specific *mCat-Pfkfb3*-expressing mice

To do this, we first mated *mCat^lox^/+*^20^ with *Pfkfb3^lox^/+* mice, which generated the double transgenic *Pfkfb3^lox^/+;mCat^lox^/+* mice. Then, *Pfkfb3^lox^/+;mCat^lox^/+* mice were mated with *CamkIIα-Cre* to generate *CamkIIα-Pfkfb3-mCat* mice.

### Neurons in primary culture

Neurons in primary culture were prepared from the cortex of E14.5 embryos ^20^ from the genotypes *Pfkfb3^lox^/+*, *mCat^lox^/+* and *Pfkfb3^lox^/+;mCat^lox^/+*. Littermates were genotyped and used individually. Cell suspensions were seeded at 2.0 x 10^5^ cells/cm^2^ in poly-D-lysine (10 μg/ml) coated plastic plates in Neurobasal A supplemented with 2 mM glutamine, 5.5 mM glucose, 0.22 mM pyruvate, and 2% antioxidant (AO) B27 supplement. Cells were incubated at 37°C in a humidified 5% CO_2_-containing atmosphere. At 72 h after plating, the medium was replaced by 2% of the minus antioxidant (MAO; i.e., lacking vitamin E, vitamin E acetate, superoxide dismutase, catalase and glutathione) B27 supplement and they were transduced with AAV-CMV-Cre-GFP or AAV-CMV-GFP to obtain transgene expression or controls, as indicated, and further incubated with fresh medium until day 7. Immunocytochemical analysis revealed that ∼99.5% of cells were neurons^35^.

### Genotyping by polymerase chain reaction (PCR)

For *Pfkfb3^lox^/+* genotyping, a PCR with the following primers was performed 5’-CGTGATCTGCAACTCCAGTCTTTC-3’; 5’-CCCAGATGACTACTTATCCTCCCA and 5’-TCCCAGTCATAGCTGTCCCTCTTC-3’. PCR conditions were 5 min at 98 °C, 25 cycles of 5 s at 98 °C, 5 s at 60 °C, 20 s at 72 °C and final extension of 2 min at 72 °Cresulting in a 82-bp band for *Pfkfb3^lox/lox^* mice and 217-bp for wild type. Primers for genotyping the mCAT allele were 5’-CTCCCAAAGTCGCTCTGAGTTGTTATCA-3’, 5’-CGATTTGTGGTGTATGTAACTAATCTGTCTGG-3’ and 5’-GCAGTGAGAAGAGTACCACCATGAGTCC-3’, which yielded a 778-bp band for the wild-type allele and a 245-bp band for the mCAT allele.PCR conditions were 5 min at 94 °C, 35 cycles of 30 s at 94°C, 30 s at 65°C, 3 min at 68 °C and 8 min at 68 °C. PCR products were resolved in 3% agarose gel using the 1 Kb DNA ladder plus (Thermo Fisher Scientific).

### Immunomagnetic purification of neurons and astrocytes from adult brain

The brain (minus cerebellum and olfactory bulb) was dissociated using the adult mouse brain dissociation kit (Miltenyi Biotec). The tissue, once clean, was fragmented with a sterile scalpel in 2 ml per hemisphere of a disintegration solution (Earle’s Balanced Salt Solution, EBSS, 116 mM NaCl, 5.4 mM KCl, 1.5 mM MgSO_4_, 26 mM NaHCO_3_, 1.01 mM NaH_2_PO_4_•2H_2_O, 4 mM glucose, 10 mg/l phenol red, supplemented with 14.4 μl/ml albumin and 26 μl/ml DNase type I, pH 7.2,), and it was trypsinized (10.8 μl/ml trypsin,) at 37°C in a thermostated bath for 5 minutes, shaking frequently to avoid decantation of the tissue. Then, the suspension was triturated using a 5-ml serological pipette (5 times) and further incubated for 10 minutes under frequent shaking. Trypsin activity was stopped by adding 10% fetal calf serum, before centrifuging the tissue at 700 *g* for 5 minutes in a microfuge at 4 °C. Once the enzymatically disintegrated tissue was decanted, the pellet was resuspended in a trypsin-free disintegration solution (EBSS + 13 μl/ml DNase + 20 μl/ml albumin) for trituration using a fire-polished Pasteur pipette. Approximately 5 passages were performed for a volume of 4 ml per hemisphere. The supernatant was centrifuged for 3 min at 700 *g* and the number of cells in the pellet counted. Once a homogeneous suspension of individualized adult neural cells was achieved, cell population separations were performed using MACS® Technology using either the astrocyte-specific anti-ACSA-2 Microbead Kit or the neuron-specific Neuron Isolation Kit, according to the manufacturer’s protocol (MACS® technology). We confirmed the identity of the isolated fractions by Western blotting against neuronal (Tuj1) or astrocytic (GFAP)-specific markers^35^.

### *In vivo* viral transduction

This was carried out using a validated adeno-associated virus (AAV) strategy^36^. Essentially, AAV particles of the PHP.eB capsid (serotype), known to efficiently transduce the central nervous system *via* intravenous injection^37^, expressing Cre recombinase driven by the neuron-specific hSyn promoter (PHP.eB-AAV-hSyn-Cre-GFP) were administered intravenously (50 µl aliquots of a phosphate-buffered saline solution containing 0.001% Pluronic® F-68, Sigma-Aldrich, and 5 x 10^10^ viral genomes, VG) through the retro-orbital sinus to 2 months-old *Pfkfb3^lox^/+* mice under a brief sevoflurane anesthesia (Sevorane, Abbot, Spain, at 6% for initiation followed by ∼3% for maintenance in air with supplement O_2_ and NO_2_ −0.4 and 0.8 liters/min, respectively-using a gas distribution column, Hersill H-3, Spain, and a vaporizer, (InterMed Penlons Sigma Delta, UK). We used the retro-orbital sinus intravenous route because of the higher success rate observed when compared with the tail or temporal ones^38^. To obtain controls, *Pfkfb3^lox^/+* siblings received equivalent amounts of the same AAV particles that did not harbor Cre recombinase. Mice were used from 4 weeks after AAV injections.

### Stereotaxic injections

Nine weeks-old *Pfkfb3^lox^/+* mice were anesthetized by inhalatory induction (4%) and maintained (2.5 %) with sevoflurane (Sevorane, Abbot) in a gas mixture of 70 % N_2_O, 30 % O_2_, using a gas distribution column (Hersill H-3, Spain) and a vaporizer, (InterMed Penlons Sigma Delta, UK). Mice were placed in a stereotaxic alignment system (Model 1900, David Kopf Instruments, CA, USA) and complemented with a stereomicroscope (Nikon SMZ 645, Tokyo, Japan) and a fiber optic cold light source (Schott KL1500 compact, Mainz, Germany). Injection was performed into the arcuate nucleus at the coordinates of AP=-2.00 mm; ML= +0.25 mm; DV=-5.5 mm from the bregma using a 5-μL Hamilton syringe with a (Microliter 65RN, Hamilton, NV, USA). 26 S needle (type 2 tip).Either 0.5 μl of AAV10-CamkIIα-Cre-GFP or AAV10-CamkIIα-GFP (2.75 x 10^12^ viral genomes, VG) was injected using a 5-µl Hamilton syringe at a rate of 0.25µl/min during 2 min with a mini-pump (UltraMicroPump III, World Precision Instruments, USA). The same volume of the dye Evans Blue (Sigma) was injected in the same manner to confirm the injection site. At the end of the injection, the syringe was left in place for 5 min before being slowly removed to prevent reflux. The skin was sutured, and mice were allowed to recover in a warming cabinet (Plactronic Digital, 25 x 60, JP Selecta, Barcelona, Spain).

### Mouse perfusion, immunohistochemistry and image analysis

Animals were deeply anesthetized by intraperitoneal injection of a mixture (1:4) of xylazine hydrochloride (Rompum; Bayer) and ketamine hydrochloride/chlorbutol (Imalgene; Merial) using 1 mL of the mixture per kg of body weight, and then perfused intraaortically with 0.9 % NaCl followed by 5 mL/g body weight of Somogy’s fixative [4 % (wt/vol) paraformaldehyde, 0.2 % (wt/vol) picric acid in 0.1 phosphate buffer, pH 7.4]. After perfusion, brains were dissected out sagittally in two parts and postfixed, using Somogy’s fixative, for 2 h at room temperature. Brain blocks were then rinsed successively for 10 min, 30 min, and 2 h with 0.1 M phosphate buffer solution (PBS; pH 7.4) and sequentially immersed in 10, 20, and 30 % (wt/vol) sucrose in PBS until they sank. After cryoprotection, 10, 20 and 40-mm-thick sagittal sections were obtained with a freezing-sliding cryostat (Leica; CM1950 AgProtect).

Sections were rinsed in 0.1 M PBS three times each for 10 min and then incubated with: i)1:1000 anti-NeuN (A-60; Merck Millipore), 1:300 anti-ΠIII-Tubulin (Tuj1; T2200; Sigma), 1/500 anti-GFAP (G6171; Sigma), 1/100 anti-Pfkfb3 (H00005209-M08; Novus Biologicals), 1/500 anti-Cyclin B1 (sc7393; Santa Cruz Biotechnology), 1/500 anti-Rock2 (sc398519; Santa Cruz Biotechnology) in in 0.2 % Triton X-100 (Sigma-Aldrich) and 5 % goat serum (Jackson ImmunoReseach) in 0.1 M PBS for 72 h at 4°C; ii) fluorophore-conjugated secondary antibodies (Jackson ImmunoResearch) in 0.05 % Triton X-100 and 2 % goat serum in 0.1 M PBS for 2 h room temperature^14^. After rinsing with PBS, sections were mounted with Fluoromont (Sigma) aqueous mounting medium. Confocal images were taken with a scanning laser confocal microscope (“Spinning Disk” Roper Scientific Olympus IX81) with three lasers 405, 491 and 561 nm and equipped with 63 x PL Apo oil-immersion objective for high-resolution imaging and device digital camera (Evolve; Photometrics, Tucson, USA).

The dendrite integrity in the hippocampus was assayed by analyzing the density of Tuj1-positive dendrites in three sections per animal. Fluorescence 8-bit images were acquired as z stacks and were exported into ImageJ in tiff format for processing. Images were converted to grayscale 8-bit images and brightness/contrast was adjusted using the ImageJ “auto” function. All Tuj1-positive dendrites and GFAP-positive cells were automatically delineated using the “auto setting threshold” (default method) and “dark background” functions of ImageJ. Thresholded images were subsequently quantified as percent area (area fraction) using the “analyze-measure” function, which represents the percentage of pixels in the image that have been highlighted (% area)^14^. Values are mean ± SEM from 15 measurements.

### Terminal deoxynucleotidyl transferase dUTP nick end-labeling (TUNEL) assay

TUNEL assay was performed in 20 μm brain sections, following the manufacturer’s protocol (Roche Diagnostics, Heidelberg, Germany). Brain sections, fixed as above, were preincubated in TUNEL buffer containing 1 mM CoCl2, 140 mM sodium cacodylate and 0.3 % Triton X-100 in 30 mM Tris buffer, pH 7.2, for 30 min. After incubation at 37 °C with the TUNEL reaction mixture containing terminal deoxynucleotidyl transferase (800 U/mL) and nucleotide mixture (1μM) for 90 min, sections were rinsed with PBS and counterstained with Cy3-streptavidin (Jackson ImmunoResearch Laboratories)^39^.

#### Plasma determination of lipids and hormones

For serum triglycerides (T2449, Sigma Aldrich, USA) and fatty acids (NEFA-HR, Wako Pure Chemical, Deutschland) analyses, we used 10 μl of undiluted serum; for insulin (#A05105, SpiBio, France) and leptin (#A05176, SpiBio, France) levels, the serum was diluted four times and we used 50 μl and 100 μl, respectively. We used the protocols according to the manufacturer’ instructions.

### Glucose tolerance test

This analysis was performed by injecting intraperitoneally D-glucose 2 g/kg body weight (Sigma) in mice that had been previously fasted for 16 hours. Blood was collected from tail bleeds every 30 min to 3 hour period and the amount of plasma glucose was determined using a glucometer (FreeStyle Optium Neo, Abbot). Areas under the curve values were determined using *GraphPad Prism 8* software.

### Mitochondrial ROS

Mitochondrial ROS were determined with the fluorescent probe MitoSox (Life Technologies). Cultured cells or adult brain-cell suspensions were incubated with 2 μM of MitoSox for 30 min at 37 °C in a 5% CO_2_ atmosphere in HBSS buffer (134.2 mM NaCl, 5.26 mM KCl, 0.43 mM KH_2_PO_4_, 4.09 mM NaHCO_3_, 0.33 mM Na_2_HPO_4_·2H_2_O, 5.44 mM glucose, 20 mM HEPES and 20 mM CaCl_2_·2H_2_O, pH 7.4). The cells were then washed with phosphate-buffered saline (PBS: 136 mM NaCl; 2.7 mM KCl; 7.8 mM Na_2_HPO_4_·2H_2_O; 1.7 mM KH_2_PO_4_; pH 7.4) and collected by trypsinization. MitoSox fluorescence intensity was assessed by flow cytometry (FACScalibur flow cytometer, BD Biosciences) and expressed in arbitrary units.

### H_2_O_2_ Determination

For H_2_O_2_ assessments, AmplexRed (Life Technologies) was used. Cells were trypsinized and incubated in KRPG buffer (145 mM NaCl, 5.7 mM Na_2_HPO_4_, 4.86 mM KCl, 0.54 mM CaCl_2_, 1.22 mM MgSO_4_, 5.5 mM glucose, pH 7.35) in the presence of 9.45 μM AmplexRed containing 0.1 U/mL horseradish peroxidase. Luminescence was recorded for 2 h at 30 min intervals using a Varioskan Flash (Thermo Scientific) (excitation, 538 nm; emission, 604 nm). Slopes were used for calculations of the rates of H_2_O_2_ formation.

### Mitochondrial membrane potential

The mitochondrial membrane potential (ΔΨ_m_) was assessed through two different methodological approaches: i) MitoProbe DiIC_1_(5) (50 nM, Life Technologies) by flow cytometry (FACScalibur flow cytometer, BD Biosciences) and expressed in arbitrary units. For this purpose, cultured cells or adult brain-cell suspensions were incubated with the probe for 30 min at 37°C in PBS. ΔΨ_m_ was obtained after subtraction of the potential value determined in the presence of carbonyl cyanide-4-(trifluoromethoxy) phenylhydrazone (CCCP) (10 µM, 15 min) for each sample; ii) TMRM (10nM, Sigma-Aldrich) plus cyclosporine-H (1 μM, Sigma-Aldrich) by confocal microscopy using Operetta CLS microscope (PerkinElmer). For this aim, primary neurons were seeded onto 96-well plate (PerkinElmer) and preincubated in KRPG buffer (145 mM NaCl, 5.7 mM Na_2_HPO_4_, 4.86 mM KCl, 0.54 mM CaCl_2_, 1.22 mM MgSO_4_, 5.5 mM D-glucose, pH 7.35). Furthermore, cells were loaded with the dye TMRM 10 nM plus cyclosporine-H 1 μM in the Operetta CLS microscope (30 min at 37 °C in a 5 % CO_2_ atmosphere) and confocal images were acquired a 40X, 1.4 NA objective (PerkinElmer). Mitochondrial uncoupler carbonyl cyanide-4-(trifluoromethoxy) phenylhydrazone (CCCP, 10 μM, Sigma-Aldrich) was added for 15 min as control of mitochondrial depolarization. Finally, images were analyzed using Harmony software (PerkinElmer).

### Flow cytometric analysis of apoptotic cell death

Adult brain-cell suspensions from the hippocampus or mediobasal hypothalamus were incubated with APC-conjugated annexin-V and 7-amino-actinomycin D (7-AAD) (Becton Dickinson Biosciences) to determine quantitatively the percentage of apoptotic neurons (Syn-GFP^+^) by flow cytometry. Brain-cell suspensions were stained with annexin-V-APC and 7-AAD in binding buffer (100 mM HEPES, 140 mM NaCl, 2.5 mM CaCl_2_), according to the manufacturer’s instructions, and 10 x 10^4^ cells were analyzed, in three replicates per condition, on a FACScalibur flow cytometer (15 mW argon ion laser; CellQuest software, Becton Dickinson Biosciences), using FL4 and FL3 channels, respectively. Annexin^+^ and 7-AAD^-^ cells were considered apoptotic. The analyzer threshold was adjusted on the flow cytometer channel to exclude most of the subcellular debris to reduce the background noise owing to the neurite disruption during neuronal resuspensions. Data were expressed as percentages.

### Determination of metabolic fluxes

To assess glycolysis, pentose-phosphate pathway (PPP), and lipogenesis fluxes, we used radiometric approaches. To do this, neurons were seeded in 8 cm^2^ flasks hanging a microcentrifuge tube containing either 1 ml benzethonium hydroxide (Sigma) (for ^14^CO_2_ equilibration) or 1 ml H_2_O (for ^3^H_2_O equilibration). All incubations were carried out in KRPG (NaCl 145 mM; Na_2_HPO_4_ 5.7 mM; KCl 4.86 mM; CaCl_2_ 0.54 mM; MgSO_4_ 1.22 mM; pH 7.35) containing 5 mM D-glucose at 37 °C in the air-thermostated chamber of an orbital shaker. To ensure adequate oxygen supply for oxidative metabolism throughout the incubation period, flasks atmosphere was gassed with carbogen (5% CO_2_/95% O_2_) before sealing with a rubber cap. To measure the carbon flux from glucose to CO_2_, cells were incubated in KRPG (5 mM glucose) buffer with 0.25 µCi/ml of [6-^14^C]-or [1-^14^C]glucose^3^. Incubations were terminated after 90 mins by the addition of 0.2 ml 20% perchloric acid (Merck Millipore) and, after a further 60 mins, the tube containing benzethonium hydroxide (with the trapped ^14^CO_2_) was used to determine the radioactivity using a liquid scintillation analyzer (Tri-Carb 4810 TR, PerkinElmer). The flux of lipogenesis was measured by assaying the rate of [U-^14^C]glucose incorporation into lipids with a similar strategy using 3 μCi/ml of D-[6-^14^C]glucose in KRPG buffer (5 mM D-glucose) for 3 hours^40^. After incubations were terminated with 0.2 ml 20% perchloric acid, the cells were washed twice with PBS, recollected and centrifuged at 500 *g* for 5 min. The supernatant was discarded and the pellet was re-suspended in 500 μL of a mixture chloroform/methanol (2: 1, v/v)^41^ for 16 h at −20 °C. The extract was washed with 250 μLof 0.3 % (w/v) NaCl saturated with chloroform. The samples were centrifuged at 1,500 *g* for 15 min and the aqueous phase, containing the cellular hydrosoluble components, was discarded. Later, this same process was repeated, this time using 250 μL of 0.3 % (w/v) NaCl saturated with chloroform plus 180 μL of methanol. The resulting chloroformic phase was passed to a new tube. Every step was performed at 4 °C. An aliquot of 50 μL of the organic phase containing the lipid fraction, was used for the measurement of the radioactivity incorporated into total lipids. The glycolytic flux was measured by assaying the rate of ^3^H_2_O production from [3-^3^H]glucose using a similar strategy using 3 μCi/ml of D-[3-^3^H]glucose in KRPG buffer (5 mM D-glucose) for 120 min^3^. After incubations were terminated with 0.2 ml 20% perchloric acid, the cells were further incubated for 72 h to allow for ^3^H_2_O equilibration with H_2_O present in the central microcentrifuge tube. The ^3^H_2_O was then measured by liquid scintillation counting (Tri-Carb 4810 TR, PerkinElmer). The specific radioactivity was used for the calculations. Under these experimental conditions, 75% of the produced ^14^CO_2_ and 28% of the produced ^3^H_2_O were recovered and were taken into account for the calculations^3^.

### Lactate and glucose determinations

Lactate concentrations were measured in the culture medium spectrophotometrically^3^ by the determination of the increments in the absorbance of the samples at 340 nm in a mixture containing 1 mM NAD^+^, 8.25 U lactate dehydrogenase in 0.25 M glycine, 0.5 M hydrazine and 1 mM EDTA buffer, pH 9.5. D-glucose was measured spectrophotometrically^3^ by reading the increase in NADPH(H^+^) absorbance at 340 nm produced in two coupled reactions catalyzed by hexokinase and glucose-6-phosphate dehydrogenase (G6PD) (Roche diagnostics Corporation, Mannheim, Germany) after 10 min of incubation.

### Oxygen consumption rate assessment

Oxygen consumption rates of primary neurons were measured in real-time in an XFe24 Extracellular Flux Analyzer (Seahorse Bioscience; Seahorse Wave Desktop software 2.6.1.56). This equipment measures the extracellular medium O_2_ flux changes of cells seeded in XFe24-well plates. Regular cell medium was removed and washed twice with DMEM running medium (XF assay modified supplemented with 5 mM glucose, 2 mM L-glutamine, 1 mM sodium pyruvate, 5 mM HEPES, pH 7.4) and incubated at 37°C without CO_2_ for 30 minutes to allow cells to pre-equilibrate with the assay medium. Oligomycin, FCCP and a mixture of rotenone and antimycin, diluted in DMEM running medium were loaded into port-A, port-B and port-C, respectively. Final concentrations in XFe24 cell culture microplates were 1 μM oligomycin, 2 μM FCCP, 1 μM rotenone and 1 μM antimycin. The sequence of measurements was as follows. Basal level of oxygen consumption rate (OCR) was measured 3 times, and then port-A was injected and mixed for 3 min, after OCR was measured 3 times for 3 min. Same protocol with port-B and port-C. OCR was measured after each injection to determine mitochondrial or non-mitochondrial contribution to OCR. All measurements were normalized to average three measurements of the basal (starting) level of cellular OCR of each well. Each sample was measured in 3–5 replicas. Experiments were repeated 3 times in biologically independent culture preparations. Non-mitochondrial OCR was determined by OCR after antimycin plus rotenone injection. Maximal respiration was determined by the maximum OCR rate after FCCP injection minus non-mitochondrial OCR. ATP production was determined by the last OCR measurement before oligomycin injection minus the minimum OCR measurement after oligomycin injection.

### Activity of mitochondrial complexes

Cells were collected and suspended in PBS (pH 7.0). After three cycles of freeze/thawing, to ensure cellular disruption, complex I, complex II-III, and citrate synthase activities were determined. Rotenone-sensitive NADH-ubiquinone oxidoreductase activity (complex I)^42^ was measured in KH_2_PO_4_ (20 mM, pH 7.2) in the presence of 8 mM MgCl_2_, 2.5 mg/ml BSA, 0.15 mM NADH and 1 mM KCN. Changes in absorbance at 340 nm (30 °C) (e=6.81 mM^-1^cm^-1^) were recorded after the addition of 50 µM of ubiquinone and 10 µM of rotenone. Deactive complex I was determined after N-ethylmaleimide (NEM) treatment of cell homogenates (10 mM, 15 minutes, 15°C). Complex I activity in the presence of NEM exclusively reflects the active form of complex I, since NEM blocks the transition from de-active to active conformation. Complex II-III (succinate-cytochrome *c* oxidoreductase) activity^43^ was determined in the presence of 100 mM phosphate buffer, plus 0.6 mM EDTA(K^+^), 2 mM KCN and 200 µM of cytochrome *c*. Changes in absorbance were recorded (550 nm; 30 °C) (e =19.2 mM^-1^cm^-1^) after the addition of 20 mM of succinate and 10 µM of antimycin A. Citrate synthase activity^44^ was measured in the presence of 93 mM of Tris-HCl, 0.1 % (v/v) triton X-100, 0.2 mM acetyl-CoA, 0.2 mM DTNB; the reaction was started with 0.2 mM of oxaloacetate, and the absorbance was recorded at 412 nm (30 °C) (e =13.6 mM^-1^cm^-1^).

### Determination of total, reduced and oxidized glutathione

Cells were lysed with 1 % (wt/vol) of sulfosalicylic acid and centrifuged at 13,000 *g* for 5 min at 4 °C, and the supernatants were used for the determination of total glutathione (reduced glutathione concentration plus twice the concentration of oxidized glutathione), by using oxidized glutathione (GSSG; 0-50 μM) as a standard. Total glutathione was measured in reaction buffer (0.1 mM NaHPO_4_, 1 mM EDTA, 0.3 mM DTNB, 0.4 mM NADPH and glutathione reductase at 1 U mL^-1^, pH 7.5) by recording the increase in the absorbance at 405 nm after the reaction of reduced glutathione with 5,5’-dithiobis(2-nitrobenzoic acid) (DTNB) for 2.5 min at 15-s intervals using a Varioskan Flash (Thermo Fisher). GSSG was quantified after derivatization of GSH with 2-vinylpyridine, by using similarly treated GSSG standards (0-5 μM), and the results are expressed as the oxidized glutathione/reduced glutathione ratio (GSSG/GSH).

### NAD^+^ determinations

To determine oxidized nicotinamide adenine dinucleotide (NAD^+^) was used a bioluminescent assay NAD/NADH-Glo^TM^ Assay (Promega). Neurons were seeded onto 96-well plates and incubated 30-60 min with NAD/NADH-Glo^TM^ Detection Reagent containing reductase, reductase substrate, NAD cycling enzyme and NAD cycling substrate, according to manufacturer’s recommendations. Detergent present in the reagent lysed cells, allowing detection of total cellular NAD^+^. Due to the cycling of the coupled enzymatic reactions, the light signal will continue to increase after adding the Reagent to the sample. The luminescent signal remains proportional to the starting amount of NAD^+^ within the linear range of the assay. To measure only the oxidized (NAD^+^) form is necessary treat the cells with 25 μL of 0.4 N HCl to destroy the reduced forms (NADH). Results are expressed as mean ± SEM (pmol/mg of protein) using standard curve (0-400 nM).

### Sirtuin activity assay

To evaluate sirtuin activity that possess either histone deacetylase or mono-ribosyltransferase activity, was used the fluorimetric sirtuin activity assay kit (#K324-100; BioVision). Neurons were seeded onto 96-well plates and lysed with 300 μL of cold homogenization buffer containing protease inhibitor cocktail. Then, lysates were transferred to a cold microfuge tube and agitated on a rotary shaker at 4 °C for 15 min. Finally, 50 μL of cell homogenate was incubated at 37 °C for 30-60 min with 40 μL of reaction mix. After incubation, 10 μL of Developer was added to each sample except standards (p53-AFC substrate; 0-1,000 pmol/well) and were incubated 15 min at 37 °C. Next, fluorescence was recorded (Ex/Em= 400/505 nm) in end point mode. In this protocol the acetylated p53-AFC substrate is deacetylated by sirtuins in the presence of NAD^+^ to generate the deacetylated p53-AFC substrate, nicotinamide and O-acetyl-ADP ribose. Cleavage of the deacetylated p53-AFC substrate by the developer releases the fluorescent group (AFC; 400/505 nm). Trichostatin A is added to the reaction to specifically inhibit HDAC’s in samples.

### Fructose-2,6 bisphosphate (F26BP) determinations

For F26BP determinations, cells were lysed in 0.1 N NaOH and centrifuged (20,000 x *g*, 20 min). An aliquot of the homogenate was used for protein determination, and the remaining sample was heated at 80 °C (5 min), centrifuged (20,000 x *g*, 20 min) and the resulting supernatant used for the determination of F26BP concentrations using the coupled enzymatic reaction and F26BP standards, as previously described^45^. This approach reveals the relative abundance of F26BP generated by PFKFB3 by the coupled enzymatic activities of PFK1 (Sigma) (in the presence of 1 mM fructose-6-phosphate and 0.5 mM pyrophosphate), aldolase (Sigma), and triose-phosphate isomerase/glycerol-3-phosphate dehydrogenase (Sigma). This reaction generates glycerol-3-phosphate and oxidizes NADH (Sigma), producing a reduction in the absorbance at 340 nm that is monitored spectrophotometrically.

### Protein determinations

Protein samples were quantified by the BCA protein assay kit (Thermo) using BSA as a standard.

### Western Blotting

Cells were lysed in RIPA buffer (1% sodium dodecylsulfate, 10 mM ethylenediaminetetraacetic acid (EDTA), 1 % (vol/vol) Triton X-100, 150 mM NaCl and 10 mM Na_2_HPO_4_, pH 7.0), supplemented with protease inhibitor mixture (Sigma), 100 μM phenylmethylsulfonyl fluoride, and phosphatase inhibitors (1 mM o-vanadate). Samples were boiled for 5 min. Aliquots of cell lysates (40-60 μg of protein) were subjected to SDS/PAGE on an 8 to 15% (vol/vol) acrylamide gel (MiniProtean; Bio-Rad) including PageRuler Prestained Protein Ladder (Thermo). The resolved proteins were transferred electrophoretically to nitrocellulose membranes (0.2 µm, BioRad). Membranes were blocked with 5% (wt/vol) low-fat milk in TTBS (20 mM Tris, 150 mM NaCl, and 0.1% (vol/vol) Tween 20, pH 7.5) for 1 h. After blocking, membranes were immunoblotted with primary antibodies overnight at 4 °C. After incubation with horseradish peroxidase conjugated goat anti-rabbit IgG (1/10,000, Santa Cruz Biotechnologies), goat anti-mouse IgG (1/10,000, Bio-Rad), rabbit anti-goat IgG (1/10,000, Abcam) or goat anti-rabbit IgG (1/3,000, Bio-Rad), membranes were immediately incubated with the enhanced chemiluminescence kit WesternBright ECL (Advansta), or SuperSignal West Femto (Thermo) before exposure to Fuji Medical X-Ray film (Fujifilm), and the autoradiograms were scanned. At least three biologically independent replicates were always performed, although only one representative Western blot is shown in the main figures. The protein abundances of all Western blots per condition were measured by densitometry of the bands on the films using ImageJ 1.48u4 software (National Institutes of Health) and were normalized per the loading control protein. The resulting values were used for the statistical analysis. Uncropped scans of Western blots replicas are shown in the Source Data file.

### Primary antibodies for Western blotting

Immunoblotting was performed with anti-LC3B (1/1,000) (#2775; Cell Signaling), anti-p62 (1/1,000) (P0067; Sigma), anti-Atg7 (1/1,000) (#2631; Cell Signaling), anti-acetylated Lysine (1/1,000) (#9441; Cell Signaling), anti-Beclin (1/1,000) (#3495; Cell Signaling), anti TOMM20 (1/1,000) (ab56783; Abcam) anti-Pfkfb3 (1/500) (H00005209-M08; Novus Biologicals), anti-VDAC (1/1,000) (PC548; Calbiochem), anti-heat-shock protein-60 (HSP60) (1/1,000) (ab46798; Abcam), anti-NDUFS1 (1/500) (sc-50132; Santa Cruz Biotechnology), anti-NDUFA9 (1/1,000) (ab14713; Abcam), anti-UQCRC2 (1/1,000) (ab14745; Abcam), anti-SDHA (1/1,000) (ab14715; Abcam), anti-ATPß (1/1,000) (MS503, MitoSciences), anti-GFAP (1/500) (G6171; Sigma), anti-ß-tubulin III (1/300) (T2200; Sigma) and anti-β-actin (1/30,000) (A5441; Sigma).

### Mitochondrial isolation

To obtain the mitochondrial fraction, cell pellets were frozen at −80 °C and homogenized (ten-twelve strokes) in a glass-Teflon Potter–Elvehjem homogenizer in buffer A (83 mM sucrose and 10 mM MOPS; pH 7.2). The same volume of buffer B (250 mM sucrose and 30 mM MOPS) was added to the sample, and the homogenate was centrifuged (1,000 *g*, 5 min) to remove unbroken cells and nuclei. Centrifugation of the supernatant was then performed (12,000 *g*, 3 min) to obtain the mitochondrial fraction, which was washed in buffer C (320 mM sucrose; 1 mM EDTA and 10 mM Tris-HCl; pH 7.4) ^35^. Mitochondria were suspended in buffer D (1 M 6-aminohexanoic acid and 50 mM Bis-Tris-HCl, pH 7.0).

### Blue native gel electrophoresis

For the assessment of complex I organization, digitonin-solubilized (4 g/g) mitochondria (10–50 μg) were loaded in NativePAGE Novex 3–12% (vol/vol) gels (Life Technologies). After electrophoresis, in-gel NADH dehydrogenase activity was evaluated allowing the identification of individual complex I and complex I-containing supercomplexes bands due to the formation of purple precipitated at the location of complex I^35^. Briefly, gels were incubated in 0.1 M of Tris-HCl buffer (pH 7.4), 1 mg/ml of nitro blue tetrazolium, and 0.14 mM of NADH. Next, a direct electrotransfer was performed followed by immunoblotting against mitochondrial complex I antibodies NDUFS1 (1/500) (sc-50132; Santa Cruz Biotechnology), NDUFA9 (1/1,000) (ab14713; Abcam), complex II antibody SDHA (1/1,000) (ab14715; Abcam), complex III antibody UQCRC2 (1/1,000) (ab14745; Abcam), complex IV antibody MTCO4 (1/1,000) (ab14705; Abcam) and complex V antibody ATPß (1/1,000) (MS503, MitoSciences). Direct transfer of BNGE was performed after soaking the gels for 20 min (4 °C) in carbonate buffer (10 mM NaHCO_3_; 3 mM Na_2_CO_3_·10H_2_O; pH 9.5–10). Proteins were transferred to polyvinylidene fluoride (PVDF) membranes was carried out at 300 mA, 60 V, 1.5 h at 4 °C in carbonate buffer.

### Autophagyc flux measurement

To analyze the autophagy pathway, primary neurons or mediobasal hypothalamus (MBH) slices were incubated in the absence or presence of the inhibitors of the lysosomal proteolysis, leupeptin (100 mM) and ammonium chloride (20 mM) for 2 h. Cells or tissues were lysed and immunoblotted against LC3-II, p62, beclin1 to assess autophagy, and against TOMM20 and Vdac to assess mitophagic flux.

### Metabolomics analysis

One hemisphere from 5-6-month-old CamkIIα-Pfkfb3 mice or WT mice (n=6 for each condition), was snap frozen in liquid nitrogen and used for untargeted metabolomic analysis (Metabolon Incorporated). Ultrahigh Performance Liquid Chromatography-Tandem Mass Spectroscopy (UPLC-MS/MS) analysis detected 567 metabolites. Raw data were extracted, peak-identified, quality-control processed, curated, and normalized by the Metabolon service. Peaks were quantified with the area under the curve. For studies spanning multiple days, a data normalization step was performed to correct variation resulting from instrument inter-day tuning differences. Essentially, each compound was corrected in run-day blocks by registering the medians to equal one (1.00) and normalizing each data point proportionately. Following log2 transformation and imputation of missing values, with the minimum observed value for each compound, Welch’s two-sample t tests were used to identify compounds significantly different between experimental groups. Metabolites labelled as xenobiotics were discarded, resulting in a total of 555 metabolites included in the final analysis. Given the high false discovery rate, metabolites were filtered according to several criteria other than q-value. Such criteria included (i) biological relevance given the genetic background context; (ii) inclusion in a common pathway with a highly significant compound; (iii) residing in a similar functional biochemical family with other significant compounds; or (iv) correlation with other experimental *in vivo* approaches, namely magnetic resonance spectroscopy and mass spectrometry. Graphs corresponding to statistical analysis were carried out with *GraphPad Prism 8.0* software and the online tool *MetaboAnalyst 5.0* (as of February 2023).

### AZ67 *in vivo* administration

AZ67 (Tocris) for *in vivo* usage was dissolved in 20 % (wt/vol) PEG200 in PBS to a 20 mM concentration. *Cln7^Δex^*^2^-vehicle and *Cln7^Δex^*^2^-AZ67 mice were used for this experiment. The cannula was inserted intracerebroventricularly at the age of 8 weeks and, after at least 15 days of recovery, we injected the AZ67 at the dose identified previously (1 nmol/mouse) every 24 h^10^. The duration of the experiment was determined by the presence of hindlimb clasping the *Cln7^Δex^*^2^ vehicle-treated mice, being this 2.5 months. Weight gain of the animals was monitored during this period.

### Behavioral tests

Mice (3 or 8 months old) were left to acclimatize in the room for not less than 30 min at the same time slot of the day (2 pm-8 pm). Tracking was carried out once at a time and carefully cleaning the apparatus with 70% ethanol between trials to remove any odor cues. An ANY-box® core was used, which contained a light grey base and an adjustable perpendicular stick holding a camera and an infrared photo-beam array to track the animal movement and to detect rearing behavior, respectively. Mouse movements were tracked with the ANY-maze® software and the ANY-maze® interface to register all parameters described subsequently. For the Open Field test, a 40 cm x 40 cm x 35 cm (w, d, h) black infrared transparent Perspex insert was used, and the arena was divided in three zones, namely border (8 cm wide), center (16 % of total arena) and intermediate (the remaining area). The test lasted for 10 min, the distance travelled and the time spent in each zone were measured.

Rotarod test (Rotarod apparatus, Model 47600, Ugo Basile) was used to analyze motor balance and coordination. Mice were previously trained during three consecutive days before testing to stablish the animal’s baseline. When performing the test, the animals are placed in the cylinder under a constant acceleration of 7.2 r.p.m. When the test starts, the cylinder starts accelerating until it reaches 40 r.p.m. after 5 min. The time it takes for the animal to fall from the cylinder is recorded.

Treadmill test was used to evaluate endurance (running time) and running speed (slowness). Mice were previously acclimatized during three consecutive days (2 h/day) to allow animals to become familiar with the treadmill and to minimize psychological stress. During this period the electric grid will remain off and belt motor on but not moving. After training, animals were submitted to a graded intensity treadmill test (Model 1050 LS Exer3/6; Columbus Instruments, Columbus,OH, USA). After a warm up period the treadmill band velocity was increased until the animals were unable to run further. The initial bout of 6 min at 6 m x min^-1^ was followed by consecutive 2 m x min^-1^ increments every 2 min. Exhaustion was defined as the third time a mouse could no longer keep place with the speed of the treadmill and remained on the shock grid for 2 s rather than running. Exercise motivation was provided for all rodents by means of an electronic shock grid at the treadmill rear. However, the electric shock was used sparingly during the test.

To analyze the short-time memory, we used the Novel Object Recognition test (Stoelting) in a 40 cm x 40 cm x 35 cm (w, d, h) core with black infrared transparent Perspex insert, also tracked with the ANY-maze® software and the ANY-maze® interface to register the track of the mice. Mice were accustomed to this environment for 10 min during two consecutive days, and the test was performed on the third day. Mice were left to explore two identical equidistant cubes for 5 min (the familiarization phase) and returned for 15 min into their cage. One cube was substituted for a similar size and color sphere and mice were returned to the arena to explore the objects for other 5 min (the test phase). To score zone entries that consider the exploration of an object we consider the size of the object (3.8×3.8 cm) and the surrounding perimeter (6×6cm). The ability to recognize the sphere as a novel object was determined as discrimination index (DI) calculated as [DI = (T_N_ – T_F_) / (T_N_ + T_F_)], where T_N_ is the time spent exploring the new object (sphere) and T_F_ is the time spent exploring the familiar object (cube).

### Magnetic Resonance Spectroscopy

Localized [^1^H]MRS was performed at 11.7 Tesla using a 117/16 USR Bruker Biospec system (Bruker Biospin GmbH, Ettlinglen, Germany) interfaced to an advance III console and operating ParaVicion 6.1 under topspin software (Bruker Biospin). After fine tuning and shimming of the system water signal FWHM values typically in the 15-25 Hz range were achieved. Scanning started with the acquisition of three scout images (one coronal, one transverse and one sagittal) using a 2D-multiplane T2W RARE pulse sequence with Bruker’s default parameters. Those images were used to place the spectroscopy voxel of size 1.5 x 1.5 x 2 mm^3^ located at the right striatum of the mouse brain or 2 x 0.8 x 2 mm^3^ located in cortex (at the mid-line of the brain), always with care not to include the ventricles in the voxel (the geometry of the voxel was slightly altered to avoid this event, when necessary). At least two ^1^H-MRS spectra were acquired per scanning session per animal (5 months-old animals). The voxel was repositioned, and shimming adjustments were repeated between acquired spectra, when the spectral resolution of the obtained ^1^H-spectrum was not good. For ^1^H-MR a water suppressed PRESS sequence was used with the following parameters: Echo time = 17.336 ms (TE1 = TE2 = 8.668 ms); Repetition time = 2500 ms; Naverages = 256; Acquisition size = 2048 points; spectral width = 11 ppm (5498.53 Hz). MR spectra were fitted and quantified using LC-Model 6.3-1R^46^.

### Metabolite extraction for LC/MS analysis

Metabolites were extracted from plasma samples (thawed on ice) by adding 1 mL of cold MeOH/H_2_O (4:1) to 20µL of plasma. After thorough mixing and incubation at −20°C for 15min, samples were centrifuged (16000g, 15min, 4°C). The supernatant was recovered, dried using the miVac Concentrator (Genevac Inc, New York, U.S.A.) and resuspended into 150µL −20°C acetonitrile/water (4:1) + 15 mM ammonium acetate. The extract was transferred to LC– MS autosampler vials for analysis. Frozen brains were first freeze-dried and ground to powder using a Mixer Mill MM 400 (Retsch GmBH, Haan, Germany) operated with dry ice. An amount of powder corresponding to 20mg of fresh brain tissue was then mixed with −20°C methanol/acetonitrile/water (2:2:1) solution containing 0.1% formic acid, followed by vortexing for 1 min, incubation at −20°C for 15 min, and centrifugation (@16000g, 15min, @4°C). The supernatant was recovered, dried, and resuspended into 150µL −20°C acetonitrile/water (4:1) + 15 mM ammonium acetate. The extract was transferred to LC–MS autosampler vials for analysis.

### LC/MS Analysis

LC/MS analyses were performed using an UHPLC Vanquish FLEX chromatographic system coupled to an Orbitrap Q Exactive+ mass spectrometer (Thermo Fisher Scientific, Waltham, MA, USA) operated in negative (ESI−) or positive (ESI+) electrospray ionization modes. Metabolites were separated on a P120 HILIC-Z (2,1 × 150 mm i.d., 2,7 um) column (Agilent) at 30°C. The HILIC solvents were A, acetonitrile/H_2_0 (9:1) + 15 mM ammonium acetate and B, acetonitrile/H_2_0 (1:9) + 15 mM ammonium acetate. HILIC separation was performed at 250 μL/min with the following gradient (minutes, %B): 0, 15%; 4, 25%; 5.5, 30%; 13.5, 35%; 15.9, 50%. The column was then equilibrated for 10 min at the initial conditions before the next sample was analysed. The injection volume was 2-5 μL. MS analyses were performed in full-scan mode at a resolution of 70 000 (at 400 m/z) over the m/z range 60 – 1000. Data were acquired with the following source parameters: the capillary temperature was 250°C, the source heater temperature, 350 °C, the sheath gas flow rate, 45 a.u. (arbitrary units), the auxiliary gas flow rate, 10 a.u., the sweep gas flow rate, 1.0 a.u., the S-Lens RF level, 55 %, and the source voltage, 2.70 kV (ESI-mode) or 3.50 kV (ESI+ mode). The data were acquired in a single analytical batch. Biological samples were randomized in the analytical run and control quality (QC) samples, consisting in mix of all biological samples of the same type-i.e., plasmas and brain samples, respectively-, were injected at regular intervals throughout the experiment. Metabolites were identified by extracting the exact mass with a tolerance of 5 ppm. The raw MS isotopic profiles of metabolites were then quantified using Tracefinder (Thermo Fisher Scientific, Waltham, MA, USA). The isotopologue fractions were obtained after correcting for natural isotopic abundances using IsoCor^47,48^ (https://github.com/MetaSys-LISBP/IsoCor). The molecular ^13^C enrichments were calculated from the sum of the relative abundances of all isotopologues of a metabolite weighted by the number of ^13^C atoms in each isotopologue.

### Statistical Analysis

For simple comparisons, we used unpaired two tailed Student’s *t* test. For other multiple-values comparisons, we used one-way or two-way ANOVA followed by Tukey or Bonferroni post-hoc tests. All tests used are indicated in each figure legend. The statistical analysis was performed using the GraphPad Prism v8 software. The number of biologically independent culture preparations or animals used per experiment are indicated in the figure legends.

### Data Availability

Source data for each figure are provided with this paper: the source data and original uncropped immunoblots that support all the figures and findings of this study are available in the Source Data file.

## Acknowledgements

We acknowledge the technical assistance of Monica Resch, Lucia Martin, Estefania Prieto-Garcia and Monica Carabias-Carrasco, from the University of Salamanca. Staff members (Amandine Rocher, Hanna Kulyk, Nina Lager-Lachaud, Lara Gales, Floriant Bellvert) of MetaToul (Toulouse, France) are gratefully acknowledged for technical support and access to mass spectrometry facilities. This work was funded by the European Regional Development Fund, Agencia Estatal de Investigación (PID2019-105699RB-I00 and PID2022-138813OB-I00, to JPB), HORIZON-MSCA-2021-DN-01 (ETERNITY, 101072759 to JPB and AA), Fundación La Caixa (HR23-00793 to JPB), Instituto de Salud Carlos III (CB16/10/00282 to JPB; PI18/00285; RD16/0019/0018 to AA), Junta de Castilla y León (CS/151P20 and Escalera de Excelencia CLU-2017-03 to JPB and AA). PC is supported by grants from Methusaleum funding (Flemish government) and the Fund for Scientific Research-Flanders (FWO-Vlaanderen G0E4419N). MetaToul is part of the national infrastructure MetaboHUB, and is funded by the French National Research Agency (ANR) with grant number MetaboHUB-ANR-11-INBS-0010.

## Authors contributions statement

Conceived the idea: JPB Designed research: JPB, DJB Performed research: DJB, JA, RL, MGM, VBJ, DGR, IMR, EF, YJ, SK, JCP, DP, PRC Analyzed data: DJB, JA, RL, MGM, VBJ, DGR, IMR, EF, YJ, SK, JCP, DP, PRC, AA Contributed materials: PC Wrote the manuscript: JPB Edited and approved the manuscript: All co-authors

---

**Fig. S1.**
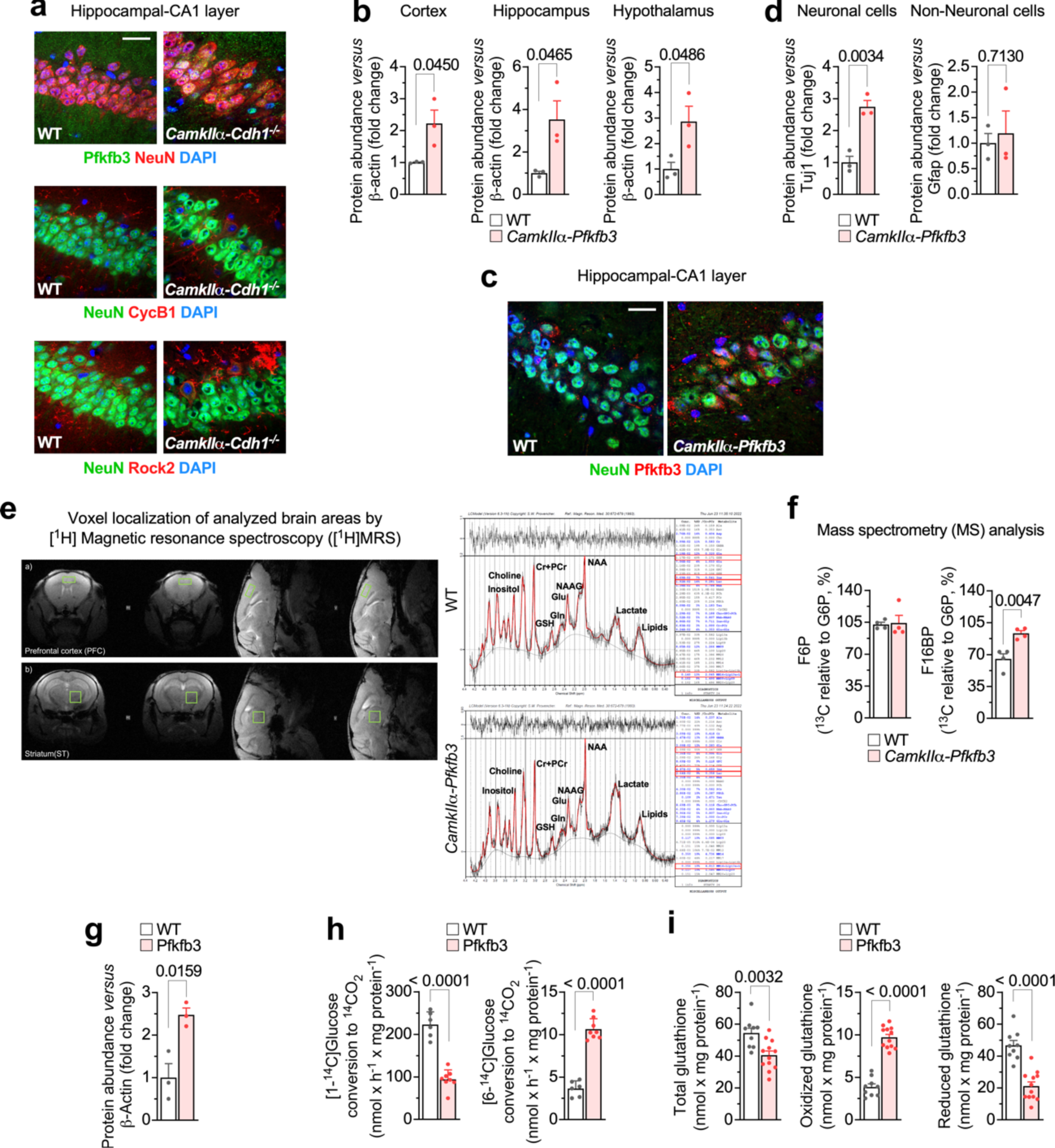
*In vivo* neuron-specific *Pfkfb3* expression activates glycolysis and inhibits PPP causing redox stress. (a) Immunocytochemical confocal images of hippocampal CA1 slices against Pfkfb3, cyclin B1 (CycB1) and Rock 2 in neuron-specific Cdh1 knockout mice (*CamkIIα-Cdh1^-/-^*). NeuN and DAPI were used to identify neurons and nuclei, respectively. Bar=20 µm. (b) Western blot quantifications for Figure 1b. Data are mean ± S.E.M. *P* value is indicated (n=3 mice per genotype; Unpaired Student’s *t*-test, two-sided). (Uncropped western blot replicas are shown in the Source Data file). (c) Immunocytochemical confocal images of hippocampal CA1 slices against Pfkfb3 in *CamkIIα-Pfkfb3* mice. NeuN and DAPI were used to identify neurons and nuclei, respectively. Bar=20 µm. (d) Western blot quantifications for Figure 1c. Data are mean ± S.E.M. *P* value is indicated (n=3 mice per genotype; Unpaired Student’s *t*-test, two-sided). (Uncropped western blot replicas are shown in the Source Data file). (e) Voxel localization of [^1^H]MRS analyzed brain areas (left) and representative ^1^H spectrum (right) in WT and *CamkIIα-Pfkfb3* mice. Related to Figure 1d,l. (f) *In vivo* F6P and F16BP concentrations analyzed by MS in brain extracts from WT and *CamkIIα-Pfkfb3* mice. Data are mean ± S.E.M. *P* value is indicated (n=4 mice per genotype; Unpaired Student’s *t*-test, two-sided). Related to Figure 1e. (g) Western blot quantifications for Figure 1g. Data are mean ± S.E.M. *P* value is indicated (n=3 mice per genotype; Unpaired Student’s *t*-test, two-sided). (Uncropped western blot replicas are shown in the Source Data file). (h) ^14^CO_2_ production from [1-^14^C]glucose (left) and [6-^14^C]glucose (right) in primary neurons. Data are mean ± S.E.M. *P* value is indicated; n=6 (WT) or 8 (Pfkfb3) biologically independent cell culture preparations; Unpaired Student’s *t*-test, two-sided). Related to Figure 1j. (i) Total (left), oxidized (middle) and reduced (right) glutathione concentrations in primary neurons. Data are mean ± S.E.M. *P* value is indicated; n=9 (WT) or 12 (Pfkfb3) biologically independent cell culture preparations; Unpaired Student’s *t*-test, two-sided). Related to Figure 1k.

**Fig. S2.**
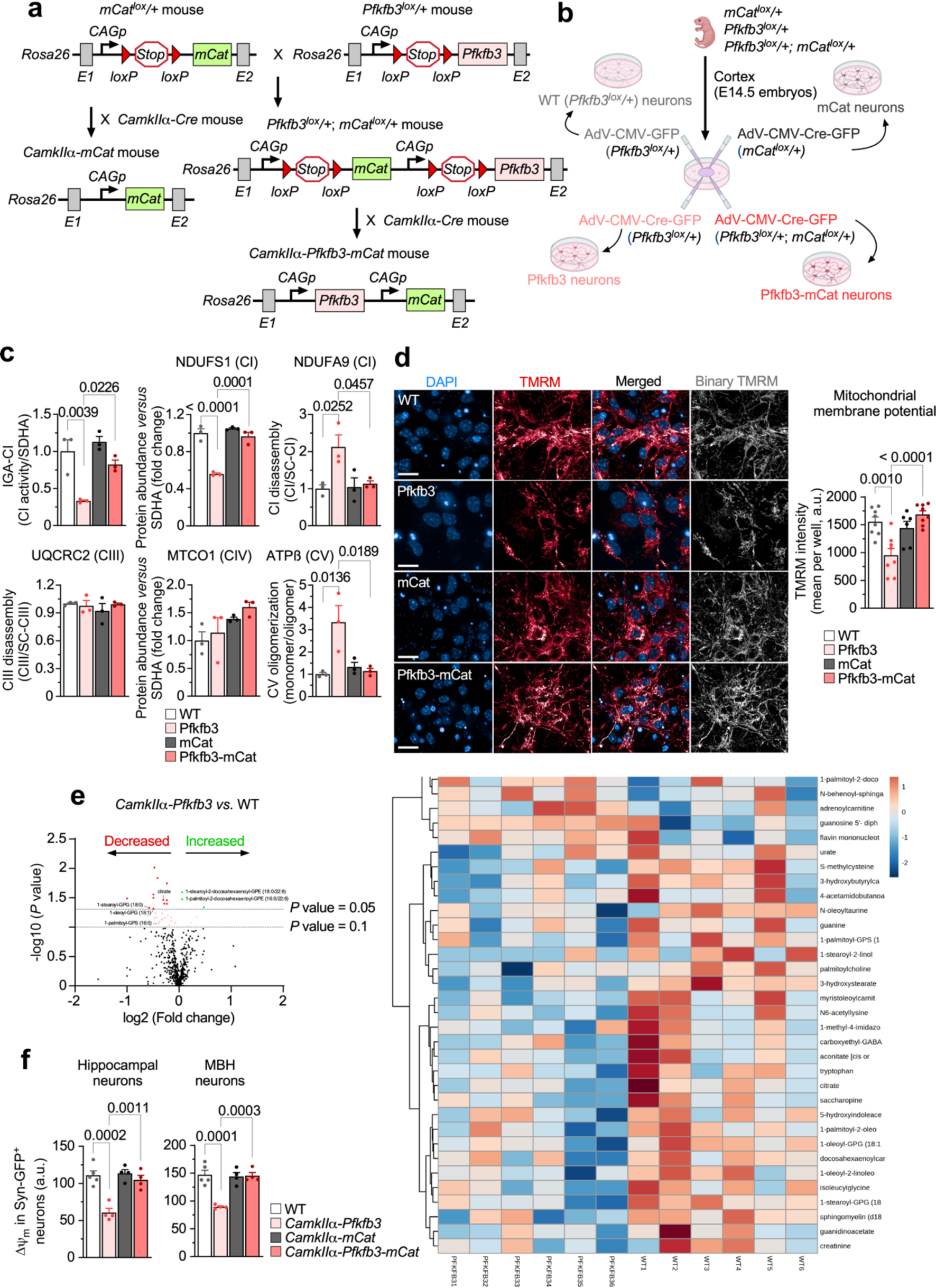
Neuron-specific *Pfkfb3* expression impairs mitochondrial bioenergetics *via* enhanced mitochondrial ROS. (a) Strategy used to generate the neuron-specific double transgenic Pfkfb3-mCat mice (*CamkIIα-Pfkfb3-mCat*) and its corresponding control (*CamkIIα-mCat*). (b) Adenoviral transduction strategy used to express double transgenic Pfkfb3-mCat in brain cortical neurons in primary culture from *Pfkfb3^lox^/+; mCat^lox^/+* mice and its corresponding control (mCat, obtained from *mCat^lox^/+*). Transduction of *Pfkfb3^lox^/+* neurons with adenovirus harboring (or not) Cre, generated Pfkfb3 or their corresponding WT (*Pfkfb3^lox^/+*) neurons. Created with BioRender.com. (c) IGA-CI and BNGE quantifications for Figure 2g. Data are mean ± S.E.M. *P* value is indicated (n=3 mice per genotype; One-way ANOVA followed by Tukey). (Uncropped western blot replicas are shown in the Source Data file). (d) ΔΨ_m_ assessment by confocal analysis in primary neurons. Data are mean ± S.E.M. *P* value is indicated; n=8 (WT, Pfkfb3, Pfkfb3-mCat) or 6 (mCat) biologically independent cell culture preparations; one-way ANOVA followed by Bonferroni. (e) Volcano plot (left) and heatmap (right) of the metabolomics analysis of the brain of *CamkIIα-Pfkfb3 versus* the corresponding WT mice. (f) ΔΨ_m_ analysis of hippocampal or MBH neurons freshly isolated from mice of the different genotypes previously transduced with adeno-associated viral particles expressing GFP under the neuron-specific *hSyn* promoter. Data are mean ± S.E.M. *P* value is indicated; n=5 (WT) or 4 (*CamkIIα-Pfkfb3*, *CamkIIα-mCat, CamkIIα-Pfkfb3-mCat*) mice; one-way ANOVA followed by Bonferroni. Related to Figure 2c.

**Fig. S3.**
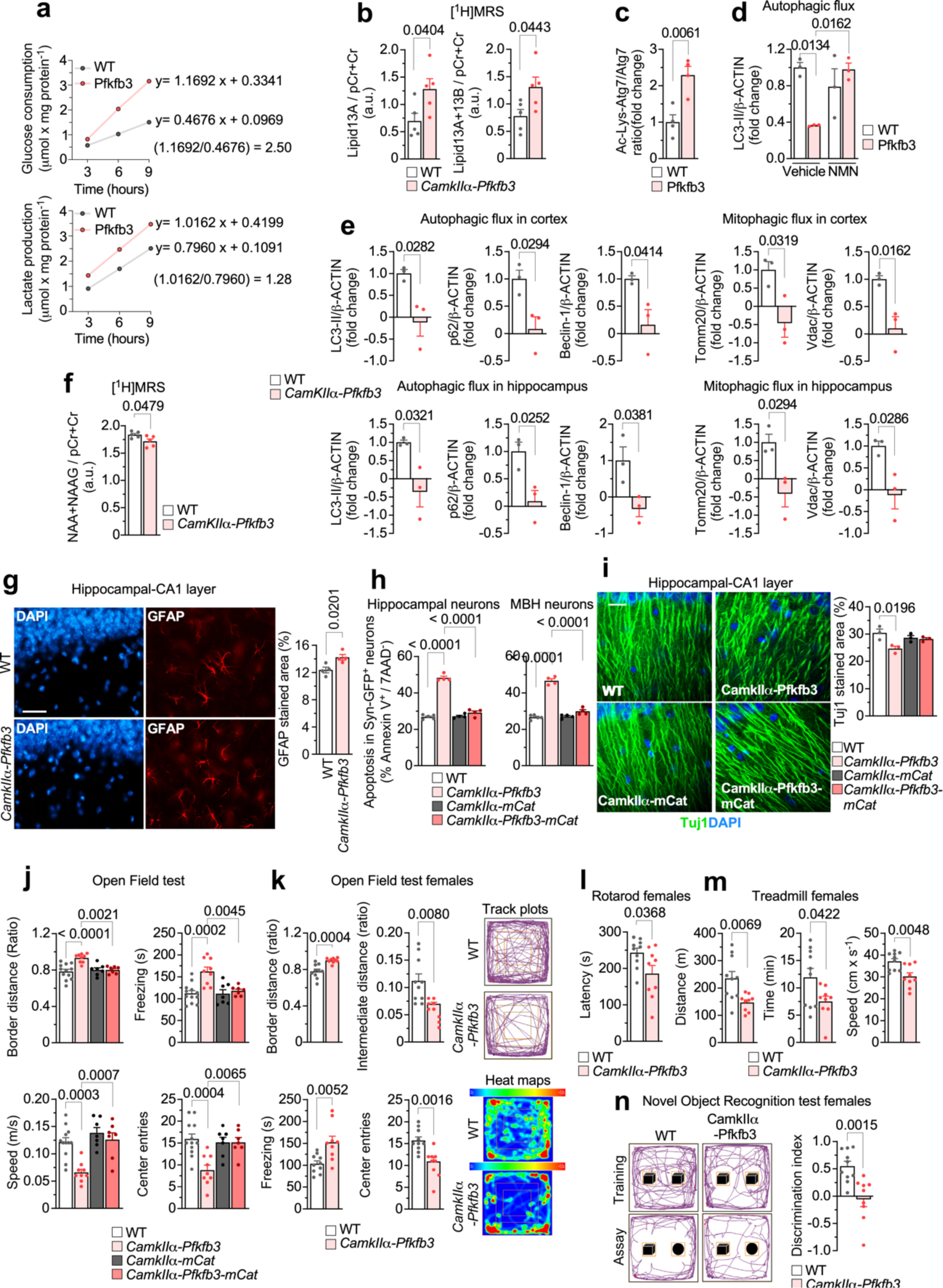
Neuron-specific *Pfkfb3* expression inhibits sirtuin-mediated autophagy causing motor and cognitive impairments *via* mitochondrial ROS. **(a)** Rates of glucose consumption (upper) and lactate release (bottom) in WT and Pfkfb3 neurons. Data are averages from 3 replica of a cell culture preparation per condition. **(b)** *In vivo* [^1^H]MRS analysis of lipid13A/(pCr+Cr) and lipid13A+13B/(pCr+Cr) in the brain of WT and *CamkIIα-Pfkfb3* mice. Data are mean ± S.E.M. *P* value is indicated; n=5 mice per genotype; Unpaired Student’s *t*-test, two-sided). **(c)** Western blot quantification for Figure 3e. Data are mean ± S.E.M. *P* value is indicated; n=4 biologically independent cell culture preparations; Unpaired Student’s *t*-test, two-sided. (Uncropped western blot replicas are shown in the Source Data file). **(d)** Western blot quantification for Figure 3f. Data are mean ± S.E.M. *P* value is indicated; n=3 biologically independent cell culture preparations; two-way ANOVA followed by Tukey. (Uncropped western blot replicas are shown in the Source Data file). **(e)** Western blot quantifications for Figure 3g. Data are mean ± S.E.M. *P* value is indicated; n=3 mice per genotype; Unpaired Student’s *t*-test, two-sided. (Uncropped western blot replicas are shown in the Source Data file). **(f)** *In vivo* [^1^H]MRS analysis of NAA+NAAG/(pCr+Cr) in the brain of WT and *CamkIIα-Pfkfb3* mice. Data are mean ± S.E.M. *P* value is indicated; n=5 mice per genotype; Unpaired Student’s *t*-test, two-sided. **(g)** Immunocytochemical analysis of hippocampal slices against GFAP and DAPI from WT and *CamkIIα-Pfkfb3* mice. Data are mean ± S.E.M. *P* value is indicated; n=4 mice per genotype; Unpaired Student’s *t*-test, two-sided. Bar=20 µm. **(h)** Apoptosis analysis of hippocampal or MBH neurons freshly isolated from mice of the different genotypes previously transduced with adeno-associated viral particles expressing GFP under the neuron-specific *hSyn* promoter. Data are mean ± S.E.M. *P* value is indicated; n=5 (WT) or 4 (*CamkIIα-Pfkfb3*, *CamkIIα-mCat, CamkIIα-Pfkfb3-mCat*) mice; one-way ANOVA followed by Bonferroni. Related to Figure 2c. **(i)** Immunocytochemical analysis of hippocampal slices against Tuj1 and DAPI from WT, *CamkIIα-Pfkfb3, CamkIIα-mCat and CamkIIα-Pfkfb3-mCat* mice. Data are mean ± S.E.M. *P* value is indicated; n=3 mice per genotype; one-way ANOVA followed by Bonferroni. Bar=20 µm. **(j)** Open field test analysis in mice. Data are mean ± S.E.M. *P* value is indicated; n=12 (WT), 9 (*CamkIIα-Pfkfb3*), 7 (*CamkIIα-mCat, CamkIIα-Pfkfb3-mCat*) mice; one-way ANOVA followed by Bonferroni. Related to Figure 3h. **(k-n)** Open field (k), rotarod (l), treadmill (m) and novel object recognition (n) tests in female mice of the indicated genotypes. Data are mean ± S.E.M. *P* value is indicated; n=10 (WT) or 9 (*CamkIIα-Pfkfb3*) mice; Unpaired Student’s *t*-test, two-sided.

**Fig. S4.**
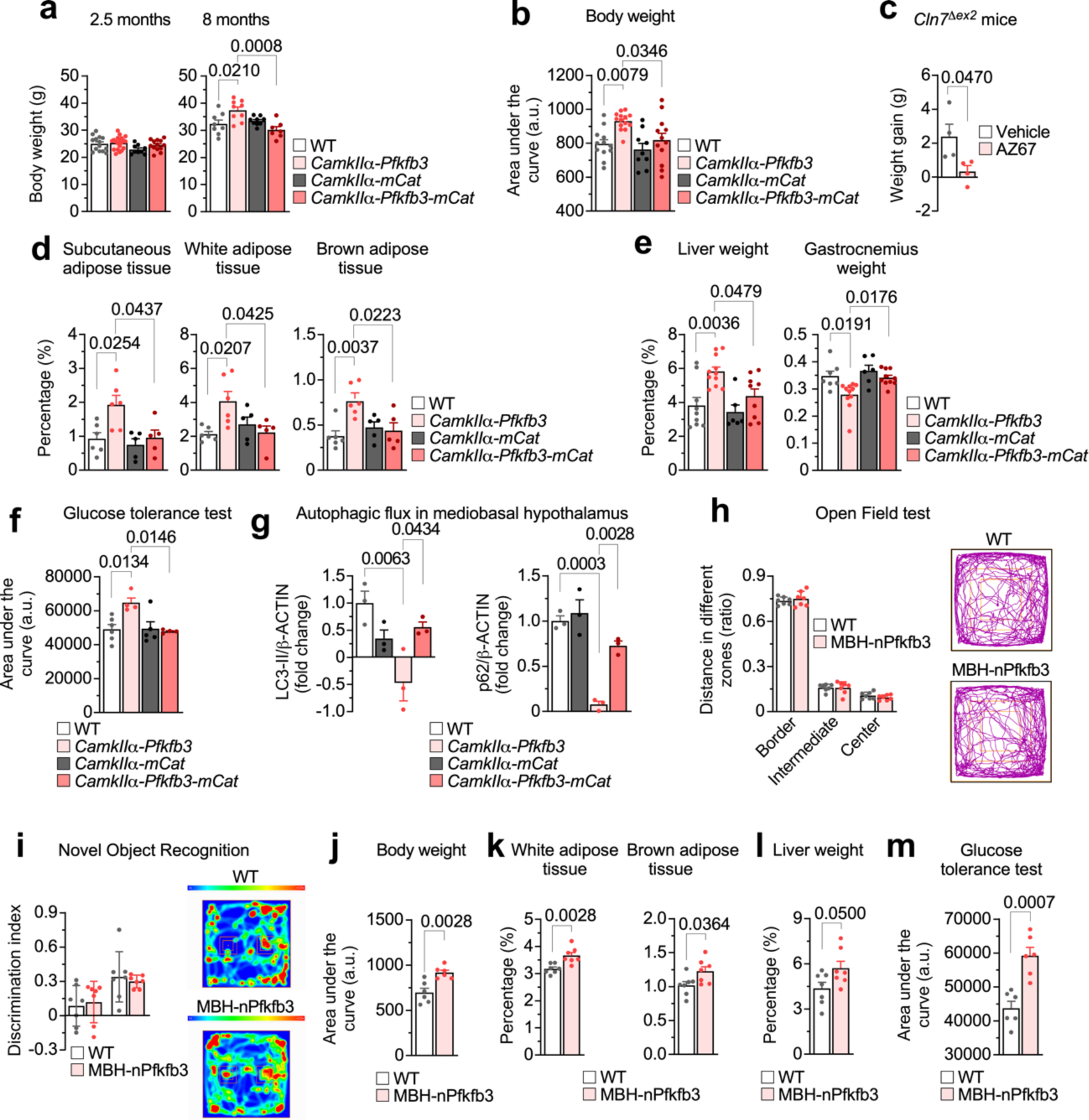
Neuronal *Pfkfb3* induces a mROS-mediated metabolic-like syndrome that is mimicked by confining Pfkfb3 expression in mediobasal hypothalamic neurons. **(a)** Body weight analysis at 2.5 and 8 months in mice. Data are mean ± S.E.M. *P* value is indicated; for 2.5 months, n=14 (WT), 20 (*CamkIIα-Pfkfb3*), 9 (*CamkIIα-mCat*) or 13 *(CamkIIα-Pfkfb3-mCat*) mice; for 8 months, n=8 (WT), 9 (*CamkIIα-Pfkfb3*, *CamkIIα-mCat*) or 7 *(CamkIIα-Pfkfb3-mCat*) mice; one-way ANOVA followed by Bonferroni. **(b)** Analysis of the area under the curve of the body weight progression of Figure 4a. Data are mean ± S.E.M. *P* value is indicated; n=12 (WT), 14 (*CamkIIα-Pfkfb3*), 9 (*CamkIIα-mCat*) or 12 *(CamkIIα-Pfkfb3-mCat*) mice; one-way ANOVA followed by Tukey. **(c)** Weight gain in *Cln7^Δex^*^2^ and WT mice that daily received the Pfkfb3 inhibitor AZ67 (1 nmol/day) (or DMSO) intracerebroventricularly for 2.5 months. Data are mean ± S.E.M. *P* value is indicated; n=4 mice per genotype; Unpaired Student’s *t*-test, two-sided. **(d)** Weight of the indicated tissues in mice. Data are mean ± S.E.M. *P* value is indicated; n=6 (WT, *CamkIIα-Pfkfb3*) or 5 (*CamkIIα-mCat*, *CamkIIα-Pfkfb3-mCat*) mice; one-way ANOVA followed by Bonferroni. **(e)** Weight of the indicated tissues in mice. Data are mean ± S.E.M. *P* value is indicated; liver, n=9 (WT), 11 (*CamkIIα-Pfkfb3*), 6 (*CamkIIα-mCat*) or 9 *(CamkIIα-Pfkfb3-mCat*) mice; gastrocnemius, n=7 (WT), 11 (*CamkIIα-Pfkfb3*), 6 (*CamkIIα-mCat*) or 10 *(CamkIIα-Pfkfb3-mCat*) mice; one-way ANOVA followed by Tukey. **(f)** Analysis of the area under the curve of glucose tolerance test of Figure 4g. Data are mean ± S.E.M. *P* value is indicated; n=6 (WT), 4 (*CamkIIα-Pfkfb3*), 5 (*CamkIIα-mCat*) or 4 (*CamkIIα-Pfkfb3-mCat*) mice; one-way ANOVA followed by Tukey. **(g)** Western blot quantifications for Figure 4 h. Data are mean ± S.E.M. *P* value is indicated; n=3 mice per genotype; one-way ANOVA followed by Tukey. (Uncropped western blot replicas are shown in the Source Data file). **(h)** Open field test analysis in WT and MBH-nPfkfb3 mice. Data are mean ± S.E.M. **(i)** Novel object recognition test analysis in WT and MBH-nPfkfb3 mice. Data are mean ± S.E.M. **(j-m)** Body weight (j), white and brown adipose tissue weight (k), liver weight (l) and area under the curve of the glucose tolerance test (m, related to Figure 4l). Data are mean ± S.E.M. *P* value is indicated; n=6 (j, m) or 7 (k, l) mice per genotype; Unpaired Student’s *t*-test, two-sided.

